# Distinct contact guidance mechanisms in single endothelial cells and in monolayers

**DOI:** 10.1101/2022.10.18.512697

**Authors:** Claire Leclech, Apoorvaa Krishnamurthy, Laurent Muller, Abdul I. Barakat

**Affiliations:** LadHyX, CNRS, Ecole Polytechnique, Institut Polytechnique de Paris, Palaiseau, France; Center for Interdisciplinary Research in Biology (CIRB), College de France, CNRS UMR7241, INSERM U1050, PSL Research University, Paris, France

**Author notes:** <u>Correspondence</u>: Prof. Abdul I. Barakat –.

**Keywords:** contact guidance, substrate topography, microgrooves, vascular endothelial cells, monolayers, basement membrane

## Abstract

In many tissues, cell shape and orientation are controlled by a combination of internal and external biophysical cues. Anisotropic substrate topography is a ubiquitous cue that leads to cellular elongation and alignment, a process termed contact guidance, whose underlying mechanisms remain incompletely understood. Additionally, whether contact guidance responses are similar in single cells and in cellular monolayers is unknown. Here, we address these questions in vascular endothelial cells (ECs) that *in vivo* form a monolayer that lines blood vessels. Culturing single ECs on microgrooved substrates that constitute an idealized mimic of anisotropic basement membrane topography elicits a strong, groove depth-dependent contact guidance response. Interestingly, this response is greatly attenuated in confluent monolayers. While contact guidance in single cells is principally driven by persistence bias of cell protrusions in the direction of the grooves and is surprisingly insensitive to actin stress fiber disruption, cell shape and alignment in dense EC monolayers are driven by the organization of the basement membrane secreted by the cells, which leads to a loss of interaction with the microgrooves. The findings of distinct contact guidance mechanisms in single ECs and in EC monolayers promise to inform strategies aimed at designing topographically patterned endovascular devices.

## Introduction

Cell shape is tightly regulated in different cell types and tissues and can exquisitely influence cell phenotype, function, and fate (Akanuma et al., 2016; Luxenburg and Zaidel-Bar, 2019; Singhvi et al., 1994). In medium and large arteries, atherosclerotic lesions initiate preferentially near branches and bifurcations where the endothelial cells (ECs) lining the blood vessels are cuboidal and randomly oriented. In contrast, zones where ECs are elongated and aligned in the direction of blood flow tend to be protected from the disease (Hahn and Schwartz, 2009). These observations suggest a prominent shape-function relationship and motivate interest in elucidating the mechanisms governing the regulation of EC morphology and alignment.

Traditionally, EC shape was thought to be driven principally by the prevailing flow characteristics within a blood vessel (Dewey et al., 1981; Helmlinger et al., 1991; Levesque et al., 1986; Lum et al., 2000). More recently, however, recognition has grown that EC shape is more broadly regulated by the integration of apical flow and substrate-derived biophysical cues acting on the cell basal surface (Dessalles et al., 2021; Liliensiek et al., 2010; Morgan et al., 2012; Song et al., 2012).

The basement membrane (BM) to which ECs adhere *in vivo* is a patterned surface that exhibits topographic features at different scales (Leclech et al., 2020). Substrate topography constitutes a biophysical basal cue that has been shown *in vitro* to influence morphology and function of various cell types (Leclech and Villard, 2020). Different types of engineered substrates have been devised to explore cellular responses to BM topography including rough surfaces, electrospun fibers, or microstructures of different shapes and sizes (Leclech and Villard, 2020). Among these systems, nano-or micro-grooved substrates can be used to mimic the anisotropy often found at different scales in the organization of various BMs, including the vascular BM (Leclech et al., 2020). Grooves have been used extensively in the literature and have been shown to elongate and align many cell types including ECs, a process known as contact guidance. Examples of contact guidance can also be found *in vivo*, most notably in the context of tumor invasion when cancer cells are able to use aligned structures in their environment to invade surrounding tissues (Sahai, 2007; Sharma et al., 2012; Weigelin et al., 2012).

Despite previous work, the mechanisms underlying contact guidance responses remain incompletely understood. One commonly invoked model posits that anisotropic topographic cues favor focal adhesion (FA) elongation in the direction of the topography while simultaneously restricting FA growth in the orthogonal direction, thereby driving the formation of aligned actin stress fibers (SFs) and the generation of anisotropic stresses that orient and elongate the cells in the direction of the topography (Ray et al., 2017). The extent to which the above model of contact guidance is universal has recently been reviewed, and evidence of possible alternative mechanisms in certain cell types has been presented (Leclech and Barakat, 2021).

The model described above centrally implicates FAs and the actin cytoskeleton in cellular contact guidance, but the role of other cytoskeletal elements remains largely unexplored. In addition, the vast majority of previous studies focused on short term contact guidance responses in single cells, while many cell types including vascular ECs are found *in vivo* as monolayers. It remains unknown whether or not the contact guidance response of cellular monolayers and its underlying mechanisms are similar to those in single cells. Several gaps thus remain in our understanding of EC shape regulation by microgrooves, which the present work aims to fill.

In this study, we use microgrooved substrates to investigate the contact guidance response of vascular ECs at densities ranging from single cells to highly confluent monolayers. We first show that when cultured on grooves, ECs become highly elongated and aligned in the groove direction and that this contact guidance response is particularly sensitive to the depth of the grooves. Remarkably, we observe a progressive loss of contact guidance with increasing cell density and time in culture. This evolution is associated with altered organization of FAs and the actin cytoskeleton. Pharmacological disruption of different cytoskeletal networks revealed that while microtubules control cell elongation at all densities, the actin cytoskeleton is surprisingly not essential in regulating the contact guidance response of single cells. We finally show that contrary to common belief, FAs are not the principal elements involved in the prominent groove depth-dependent contact guidance in single ECs. Rather, this response is driven mostly by physical constraints on cell protrusions that lead to anisotropic persistence bias of these protrusions in the direction of the grooves and consequent cell elongation. In monolayers, the progressive loss of response to topography with increasing EC density is not attributable to the emergence and reinforcement of cell-cell junctions but rather to the secretion by the cells of BM proteins whose organization dictates cell morphology and alignment.

## Results

### Groove depth is the principal determinant of endothelial cell alignment and elongation on microgrooves

In order to study the response of ECs to substrate topography, we cultured HUVECs on PDMS microstructured substrates composed of arrays of parallel grooves of different widths (w), spacing (s), and depths (d) (Fig. 1A), uniformly coated with fibronectin (Supplementary Fig. 1). Flat fibronectin-coated PDMS substrates served as controls. In line with previous reports (Antonini et al., 2015; Natale et al., 2019; Stefopoulos et al., 2017), ECs on the microgroove substrates aligned and elongated in the groove direction (Fig. 1B). It has been reported that the dimensions of grooves can modulate the extent of cell elongation and alignment (Andersson et al., 2003; Franco et al., 2011; Teixeira et al., 2003). To more systematically test this notion, we independently varied groove depth from 1 to 6 μm, width from 2 to 5 μm, and spacing from 5 to 20 μm, in each case while maintaining the other two dimensions constant. Cell orientation angle relative to the groove direction and cell circularity assessed by the ratio of the cell minor-to-major axis were analyzed using automated detection of cell boundaries based on images of monolayers stained for cell-cell junctions (VE-cadherin) shortly after attaining confluence (after 24 h of culture). While all dimensions of grooves tested led to EC elongation and alignment in the direction of the grooves, only varying the groove depth significantly changed the extent of this response (Fig. 1B,C). For instance, increasing groove depth from 1 to 6 μm (width and spacing maintained constant at 5 μm) significantly increased cell elongation and alignment in the direction of the grooves (Fig. 1C). In contrast, changing groove width from 2 to 5 μm (spacing = 5 μm, depth = 1 μm) (Fig. 1D) or groove spacing from 5 to 20 μm (width = 5 μm, depth = 1 μm) (Fig. 1E) led to much smaller effects on cell shape and alignment. Thus, for the range of dimensions studied, the contact guidance response of HUVEC monolayers is clearly more sensitive to groove depth than to either groove width or spacing. In light of these results, we focus for the rest of the study on two groove dimensions: shallow grooves (w=s=5 μm, depth = 1 μm) and deep grooves (w=s=5 μm, depth = 5 μm).

**Figure 1:**
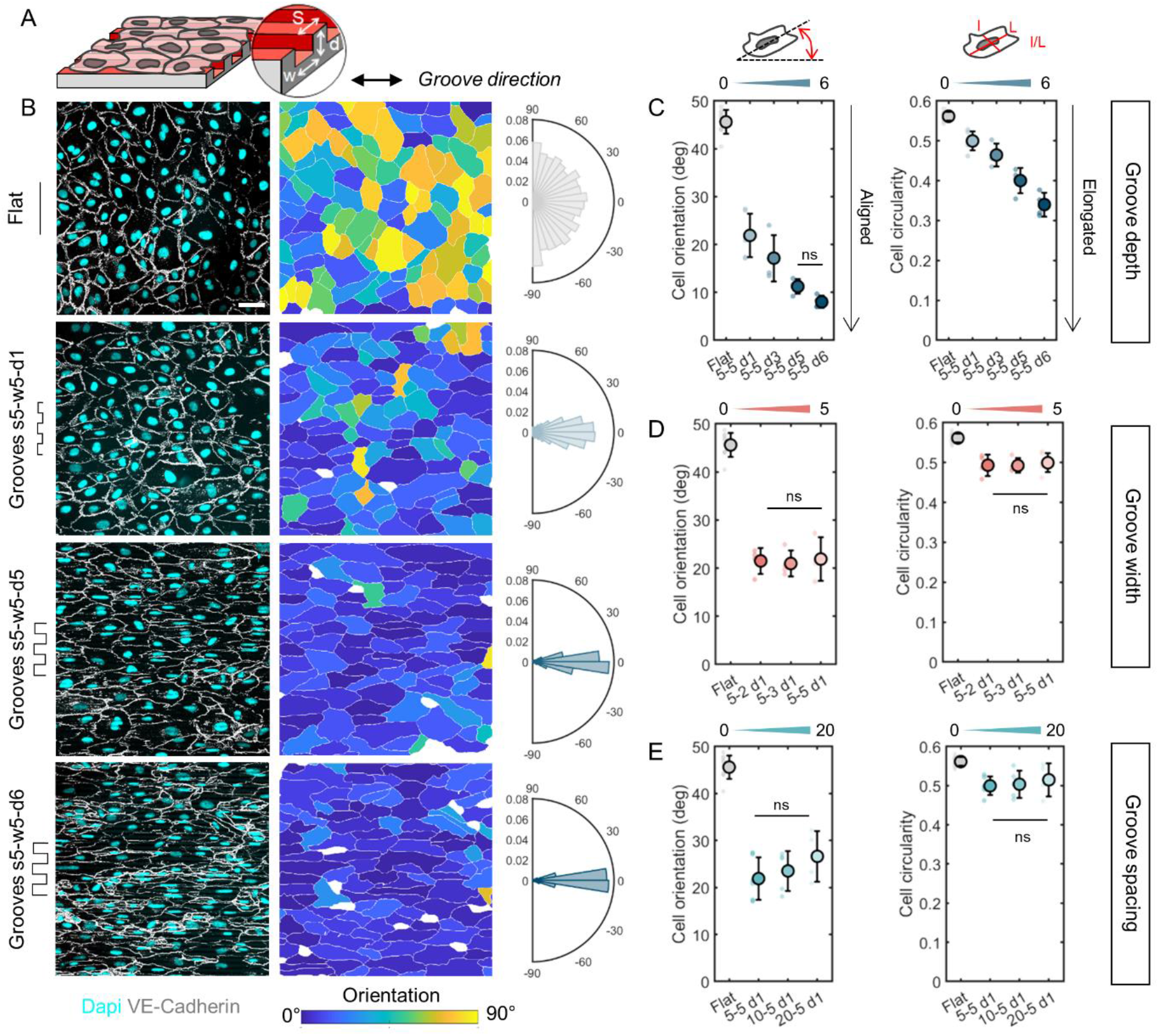
Depth-dependent contact guidance of vascular ECs on microgroove substrates. (A) Schematic of HUVEC culture on fibronectin-coated microgroove substrates of spacing s, width w, and depth d. (B) Left: immunostaining for VE-cadherin (cell-cell junctions, white) and DAPI (nucleus, cyan) of HUVEC monolayers after 24 h of culture on flat surfaces or microgrooves of width and spacing 5 μm and increasing groove depth (1, 5, and 6 μm). Scale bar, 50 μm. Center: segmented cells color-coded by orientation relative to the groove direction (0°). Right: Distributions of cell orientation angles for the different groove depths. (C) Quantification of cell orientation angle and cell circularity (ratio of minor to major axis) for different groove depths (1, 3, 5, and 6 μm), spacing = width = 5 μm. (D) Quantification of cell orientation angle and cell circularity for different groove widths (2, 3, and 5 μm), spacing= 5 μm, depth = 1 μm. (E) Quantification of cell orientation angle and cell circularity for different groove spacings (5, 10, and 20 μm), width = 5 μm, depth = 1 μm. Dots represent individual experiments. n=4 to 10 independent experiments, one-way ANOVA, Fisher’s post-test (** p < 0.001). Differences are all statistically significant unless otherwise noted (“ns” labeling). Error bars represent standard deviations.

### Endothelial cells exhibit progressive loss of response to microgrooves with increasing cell density

Most of the contact guidance studies conducted to date have been performed on single cells, which raises questions about their applicability to ECs that are found as monolayers *in vivo*, at least under normal physiological conditions. To investigate the effect of cell density on EC response to microgrooves, we analyzed EC shape on flat control surfaces as well as on 1 μm- and 5 μm-deep grooves fixed at different time points ranging from single cells at 2 h of culture to highly confluent monolayers at 72 h of culture (Fig. 2A). As expected, cell area remained stable over the first 24 h of culture (corresponding to the time point of complete surface coverage and establishment of cell-cell junctions) and subsequently decreased as cell density increased in the monolayer (Supplementary Fig. 2A). In terms of the contact guidance response to the microgrooves, cell alignment and elongation was most pronounced in single cells and then progressively decreased with time and cell density (Fig. 2B,C). Interestingly, the rate of loss of the contact guidance response with cell density (i.e. slopes of the alignment and elongation curves in Fig. 2B,C) appears to be largely similar for the two groove depths tested. It is also noteworthy that while the loss of EC elongation at the highest cell density is nearly complete with circularity reverting to values close to the controls (especially for the 1 μm-deep grooves), the loss of alignment is only partial.

**Figure 2:**
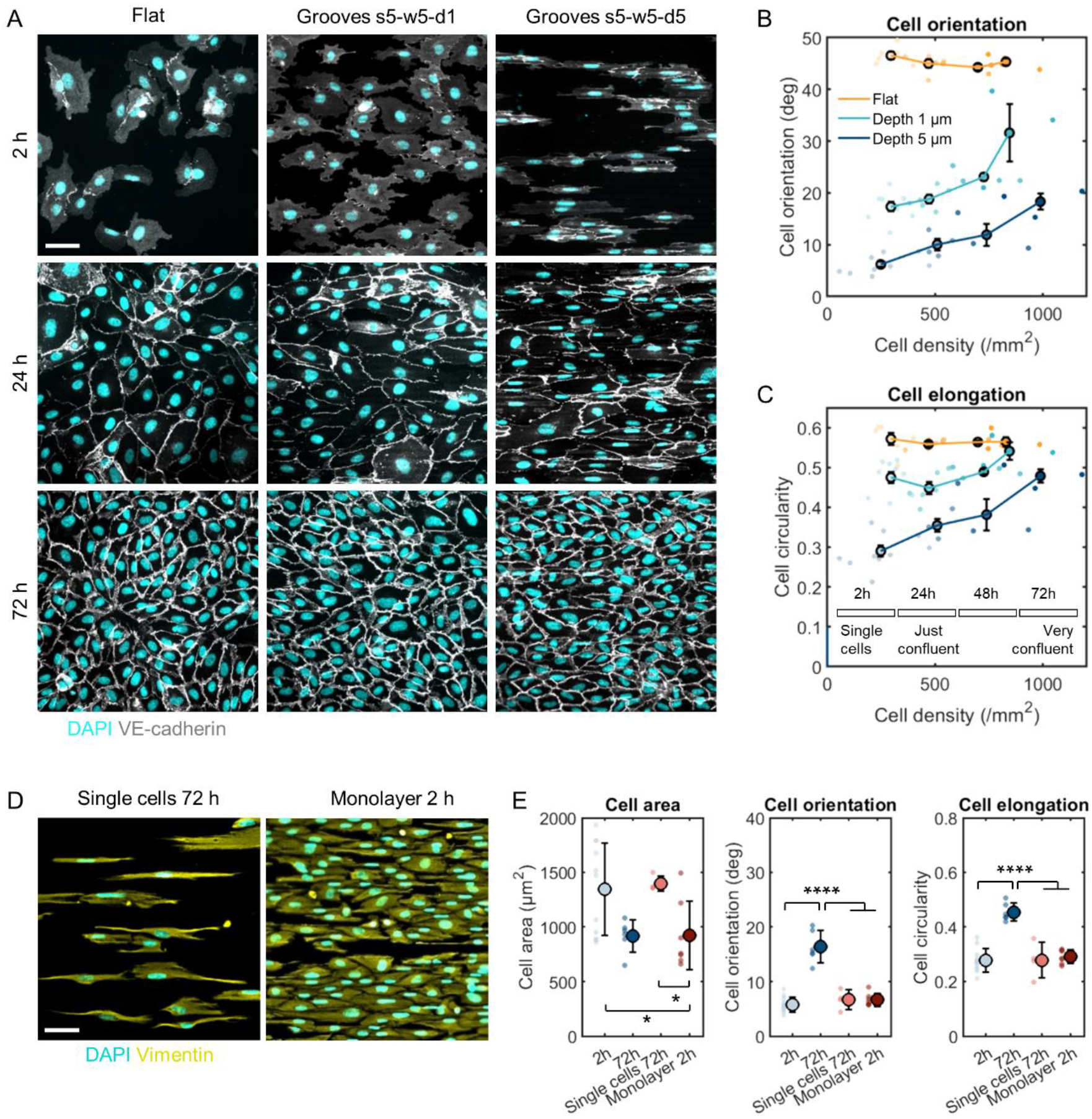
Evolution of contact guidance of ECs with time and cell density. (A) Immunostaining for VE-cadherin (white) and DAPI (cyan) of HUVECs after 2 h (single cells), 24 h (low-density monolayers), or 72 h (high-density monolayers). Scale bar, 50 μm. (B-C) Quantification of cell orientation angle and cell circularity for increasing time and cell density on flat surfaces (yellow), 1 μm-deep grooves (light blue), and 5 μm-deep grooves (dark blue). Dots represent individual experiments, regrouped by density groups corresponding to 2 h, 24 h, 48 h, and 72 h cultures, with at least 3 independent experiments per group. Error bars represent standard error of the mean (SEM). (D) Immunostaining for vimentin (yellow) and DAPI (cyan) of either single HUVECs at 72 h of culture or monolayers after 2 h of culture. Scale bar, 50 μm. (E) Quantification of cell area, cell orientation angle, and cell circularity. Dots represent individual experiments. n=4 to 11 independent experiments, one-way ANOVA, Fisher’s post-test (* p < 0.04, **** p < 0.0001). Error bars represent standard deviations.

The observations above point to a progressive loss of EC response to substrate topography with an increase in both cell density and time in culture. To discriminate between the effects of time and cell density, we cultured ECs at either a very low density so they remained as single cells even after 72 h or at very high density so as to attain a monolayer after a mere 2 h of culture (Fig. 2D). In both cases, cells exhibited prominent alignment and elongation, similar to the single cells at 2 h of culture, indicating that the reported loss of response to microgrooves necessitates both a sufficiently high cell density and a sufficiently long time in culture (Fig.2D,E).

The attenuated contact guidance in endothelial monolayers was associated with important functional ramifications. For instance, while individual ECs were polarized in the groove direction (as assessed by the position of the Golgi apparatus with respect to the nucleus), this polarization was lost in monolayers as evidenced by the Golgi being randomly localized around the nucleus (Supplementary Fig. 2B). Similarly, live-cell recordings revealed that while single ECs on microgrooves exhibited very straight trajectories that were highly constrained along the groove direction, cells within monolayers moved more freely along the axis perpendicular to the grooves (Supplementary Fig. 2C).

### Single cells and monolayers exhibit different cytoskeletal and focal adhesion organization on microgrooves

To better understand the groove depth- and cell density-dependent contact guidance responses of ECs on microgrooves, we first characterized the cytoskeletal organization in single cells and in 72 h monolayers on both 1 μm- and 5 μm-deep grooves. In single cells on microgrooves, microtubules and intermediate filaments (demarcated by staining for vimentin, the principal intermediate filament in ECs) formed a dense network at the center of the cell and around the nucleus (Fig. 3A). In contrast, actin stress fibers were organized in thick bundles localized above the grooves and spanning the entire cell (Fig. 3A and Supplementary Fig. 3). This organization was quantified by plotting the actin fluorescence intensity profile along a line through the cell in the direction orthogonal to the cell major axis, which showed periodic peaks corresponding to the bundles (red arrowheads) (Fig. 3C). Interestingly, while the microtubules and intermediate filament networks tended to follow the orientation of the cells’ major axis, the actin bundles more closely followed the groove direction. In 72 h dense monolayers, while the microtubule and intermediate filament networks broadly retained the same organization, the actin stress fibers completely relocalized to the periphery of the cell, close to cell-cell junctions, and were virtually absent from the cell center (green arrowheads) (Fig. 3B,C).

**Figure 3:**
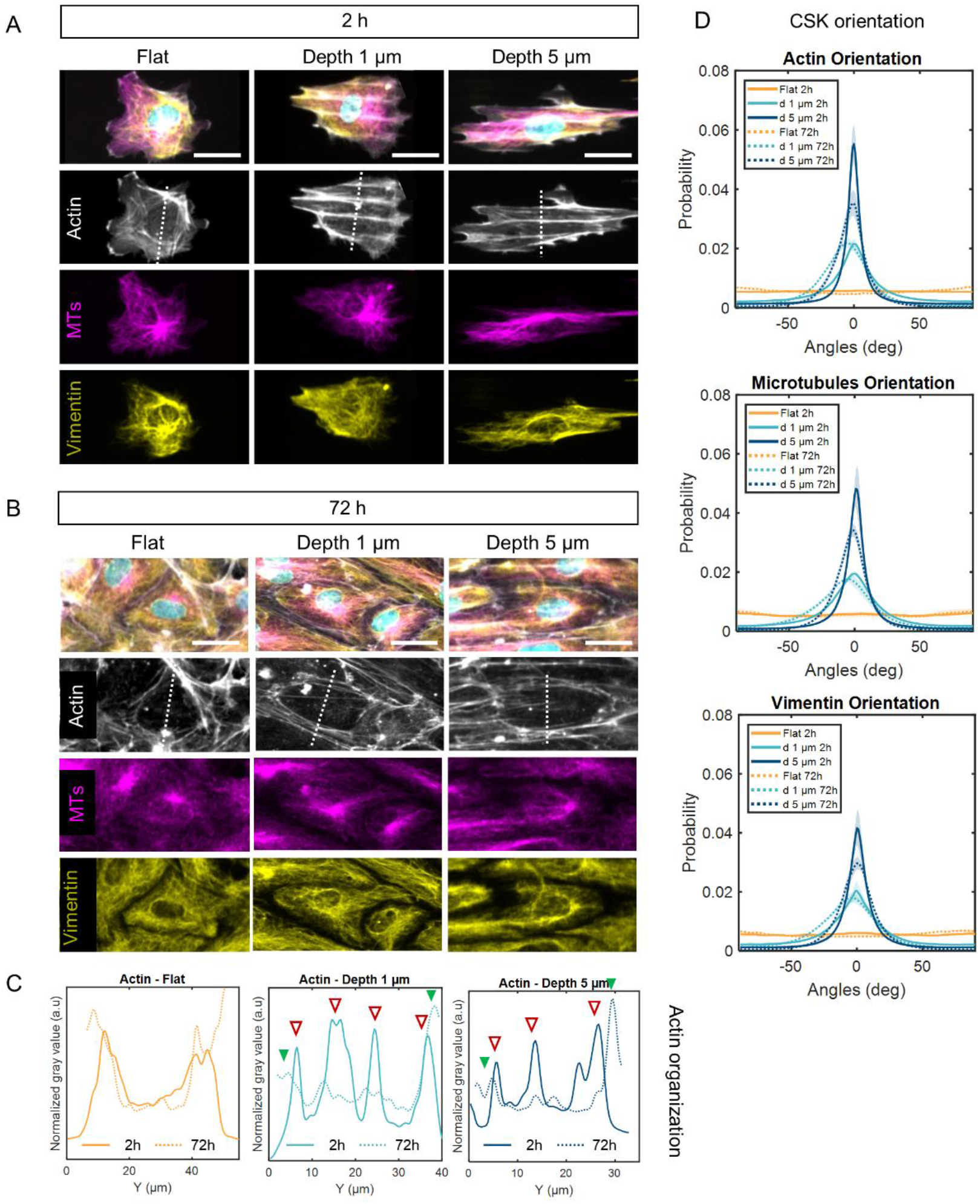
Organization of the cytoskeleton with time and groove depth. (A-B) Immunostaining for actin (phalloidin, grey), microtubules (β-tubulin, magenta), and intermediate filaments (vimentin, yellow) on flat surfaces, 1 μm-deep grooves, and 5 μm-deep grooves in single cells (2 h of culture) or monolayers (72 h of culture). Grooves are in the horizontal direction. Scale bars, 25 μm. (C) Localization of the actin cytoskeleton in the represented cells: normalized fluorescence intensity profile along the white dotted line. Arrowheads point to intensity peaks that represent bundles of actin stress fibers. (D) Orientation of the different cytoskeletal networks on flat surfaces (yellow), 1 μm-deep grooves (light blue), and 5 μm-deep grooves (dark blue) for 2 h (solid lines) or 72 h of culture (dotted lines). 0° corresponds to the groove direction.

Analysis of the orientation of the actin stress fibers, microtubules, and intermediate filaments showed that in both single cells (Fig. 3D solid lines) and monolayers (Fig. 3D dashed lines), the three cytoskeletal networks were more aligned on the deeper grooves, consistent with the more prominent cell alignment and elongation reported above (cf: Fig. 1). This difference between the two groove depths tended to be smaller in monolayers than in single cells. On 5 μm-deep grooves, the cytoskeletal networks were more aligned in single cells than in highly confluent monolayers, illustrating the loss of response to microgrooves with increased cell density. This difference was less visible for 1 μm-deep grooves. Overall, these findings are consistent with the aforementioned observations of groove depth-dependent EC alignment and elongation (especially in single cells) and the loss of response to substrate topography in high-density monolayers. In addition, the actin network appears to be the most responsive cytoskeletal element to substrate topography.

We also investigated the time evolution of the morphology and organization of FAs. In single cells, long and highly punctate FAs were visible, localizing at the ends of actin stress fibers (Fig. 4A). In contrast, a more diffuse paxillin staining was present in high-density monolayers at 72 h of culture (Fig. 4B), and mature FAs were virtually absent from the cell center and tended to localize at the cell periphery, close to the cortical actin network and cell-cell junctions. FAs were accordingly significantly smaller, less aligned, and less elongated in dense monolayers compared to those in single cells on 5 μm-deep grooves (Fig. 4C). This can be interpreted as a form of disassembly of FAs in high-density monolayers and suggests that single cells and cells in monolayers interact differently with their substrate.

**Figure 4:**
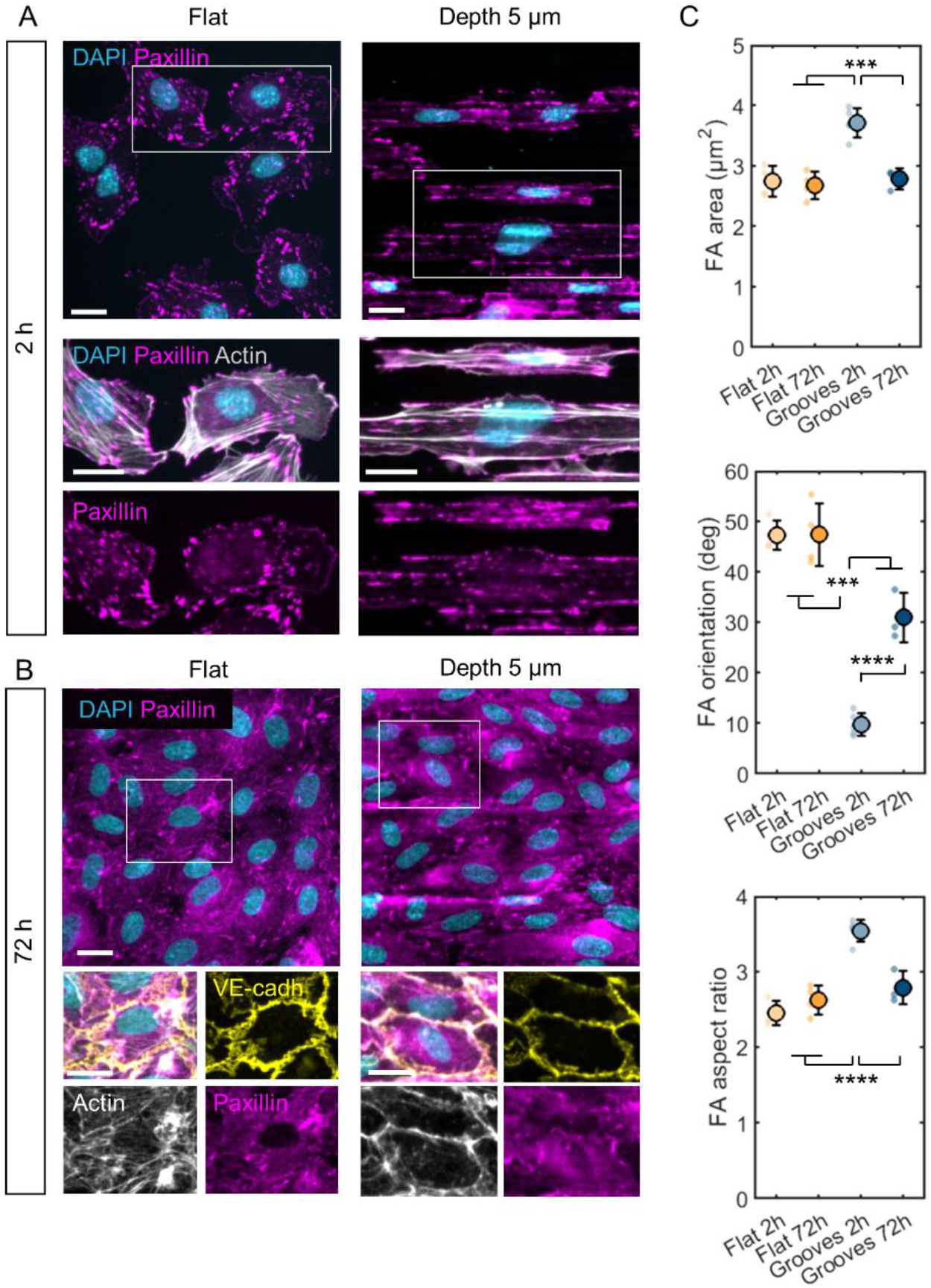
Organization of focal adhesions with time. (A-B) Immunostaining for paxillin (magenta) and DAPI (cyan) in single cells (2 h of culture) and monolayers (72 h of culture) on flat surfaces and 5 μm-deep grooves. In the zoomed-in insets, immunostaining for paxillin, actin, and DAPI (2 h) or paxillin, actin, DAPI, and VE-cadherin (yellow) (72 h of culture). Scale bars 20 μm. (C) Quantification of FA area orientation angle relative to the grooves, and aspect ratio (ratio of major to minor axis) on flat surfaces and 5 on μm-deep grooves for 2 h and 72 h. Dots represent individual experiments. n=3 to 5 independent experiments, one-way ANOVA, Fisher’s post-test (*** p < 0.001, **** p < 0.0001). Error bars represent standard deviations.

### Actin is not required for contact guidance in single ECs while microtubules drive cell elongation on microgrooves

To more functionally test the contribution of the different cytoskeletal elements to the EC contact guidance response on microgrooves, we pharmacologically disrupted cell contractility with blebbistatin (100 μM for 90 min), actin polymerization with latrunculin A (10 nM) + cytochalasin D (20 nM for 90 min), and microtubules with nocodazole (0.2 μM for 90 min) in single cells at 2 h of culture (Fig. 5A), low-density monolayers at 24 h of culture (Supplementary Fig. 4), and high-density monolayers at 72 h of culture (Fig. 5B).

**Figure 5:**
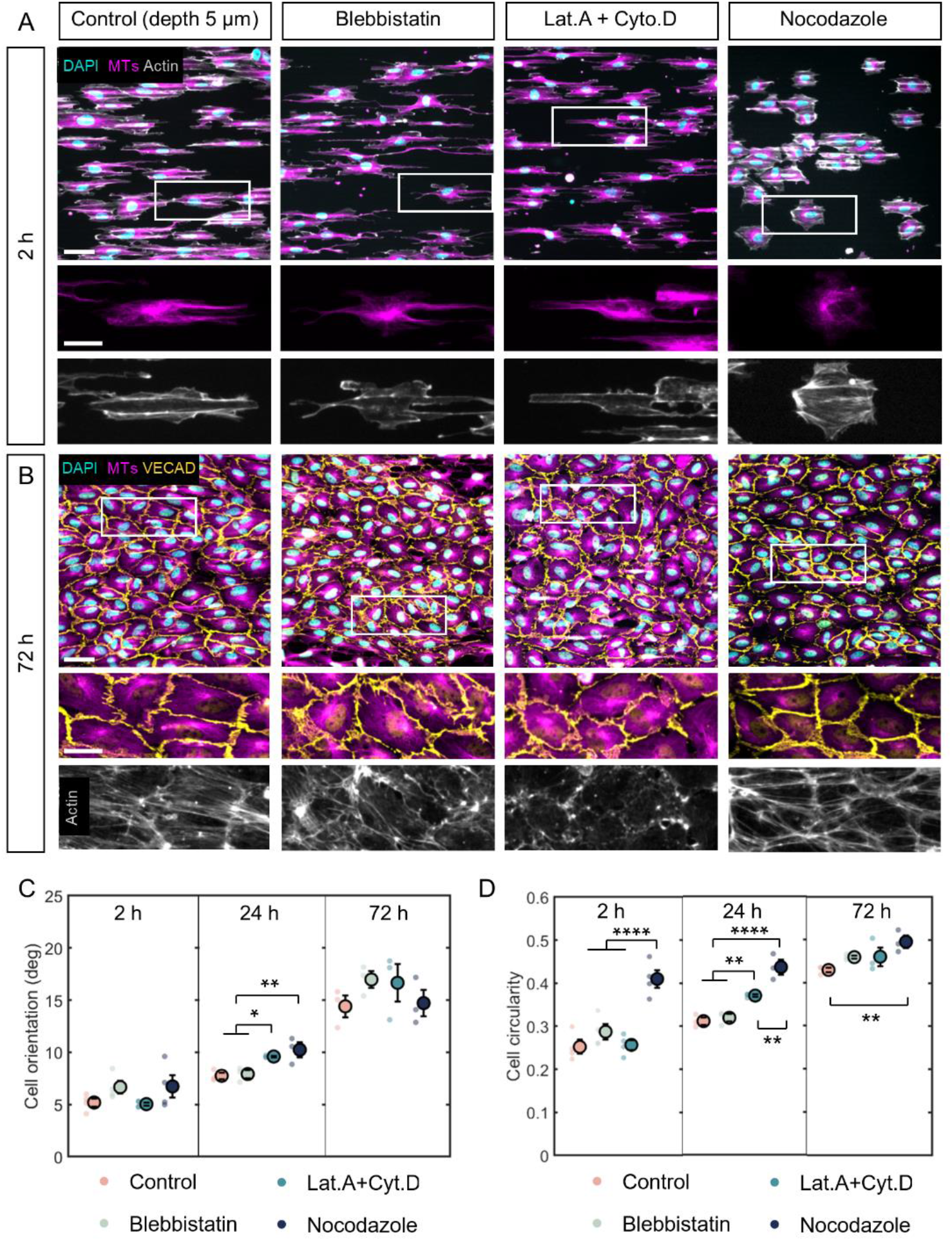
Influence of the different cytoskeletal elements on the response to microgrooves. (A-B) Immunostaining for microtubules (β-tubulin, magenta), actin (phalloidin, gray), DAPI (cyan) (2 h) and VE-cadherin (yellow, 72 h) of control cells (+DMSO) or cells treated for 90 min with 100 μM blebbistatin, 0.1 nM latrunculin A + 0.2 nM cytochalasin D, or 0.2 μM nocodazole. Scale bars, 50 μm, 25 μm (insets). Grooves are in the horizontal direction. (C-D) Quantification of cell orientation angle relative to the grooves and cell circularity (ratio of minor to major axis) upon pharmacological treatment at 2 h (single cells), 24 h (low-density monolayers), or 72 h (high-density monolayers). Dots represent individual experiments. n=3 independent experiments, one-way ANOVA, Fisher’s post-test (* p < 0.03, ** p < 0.009, **** p < 0.0001). Error bars represent standard error of the mean (SEM).

In single cells on 5 μm-deep grooves, disruption of either actin polymerization or actomyosin-mediated contractility had a visible effect on cell shape by inducing long cell protrusions (Fig. 5A) but did not change the extent of cell alignment or elongation compared to the untreated controls (Fig. 5C,D). In contrast, microtubule disruption significantly reduced cell elongation but not cell alignment. In cell monolayers, the effect of actin depolymerization became more visible, decreasing cell alignment and elongation (Fig. 5B-D). The effect of microtubule disruption on cell elongation observed in single cells persisted with increasing cell density, although the difference with the control untreated cells tended to decrease for highly confluent monolayers (Fig. 5D), likely because of the sharp decrease in elongation of the control cells at high density (cf: Fig. 2). More generally, the most significant effect of cytoskeletal disruption appears to be on cell elongation rather than on cell alignment, which suggests that these two responses may be mediated by distinct mechanisms. We also note that while microtubule depolymerization had no visible effect on cell-cell junctions, blebbistatin or latrunculin A + cytochalasin D treatment led to more wavy junctions, consistent with the documented physical interactions of actin filaments with cell-cell junctions in different cell types including ECs (Hoelzle and Svitkina, 2012).

The cytoskeletal disruption experiments described above suggest show that while disruption of either the actin or microtubule cytoskeleton does not completely abolish cell alignment and elongation in response to microgrooves, microtubules appear to be the principal elements controlling cell elongation at all cell densities. In single cells, the actin network appears to be non-essential in the contact guidance response, which raises questions about the central role of actin often evoked in the literature (Ray et al., 2017). The actin network appears however to play a more important role in confluent monolayers, potentially through cell-cell junctions with which it closely associates. Overall, these results suggest that the different cytoskeletal elements may play different roles at different cell densities.

More generally, our findings thus far point toward different regimes and potentially different mechanisms of response to topography in single cells, which are particularly sensitive to groove depth, and in monolayers where the response to topography is progressively lost. We next investigated the underlying mechanisms of these different regimes of response.

### Groove depth-dependent response in single cells relies primarily on cell protrusion guidance

As already demonstrated above, groove depth detection is a very early response of single ECs to topography, visible after only 2 h of culture (cf: Fig. 2). We next turned our attention to the mechanisms underlying this response. Because lateral constraints on FA growth imposed by the grooves have been proposed to drive cellular contact guidance (Ray et al., 2017), we first revisited the morphology and organization of FAs in single cells cultured on 1 μm-or 5 μm-deep grooves. As described already (cf: Fig. 4), FAs in single ECs exhibited long and highly punctate FAs, which were found both at the groove tops (ridges) and bottoms but localized preferentially along the edges of the ridges (Fig. 6A). Analysis of FA morphology revealed that FA area, alignment in the groove direction, and extent of elongation increased progressively with groove depth (Fig. 6B), showing that FAs are indeed responsive to the grooves and their depth.

**Figure 6:**
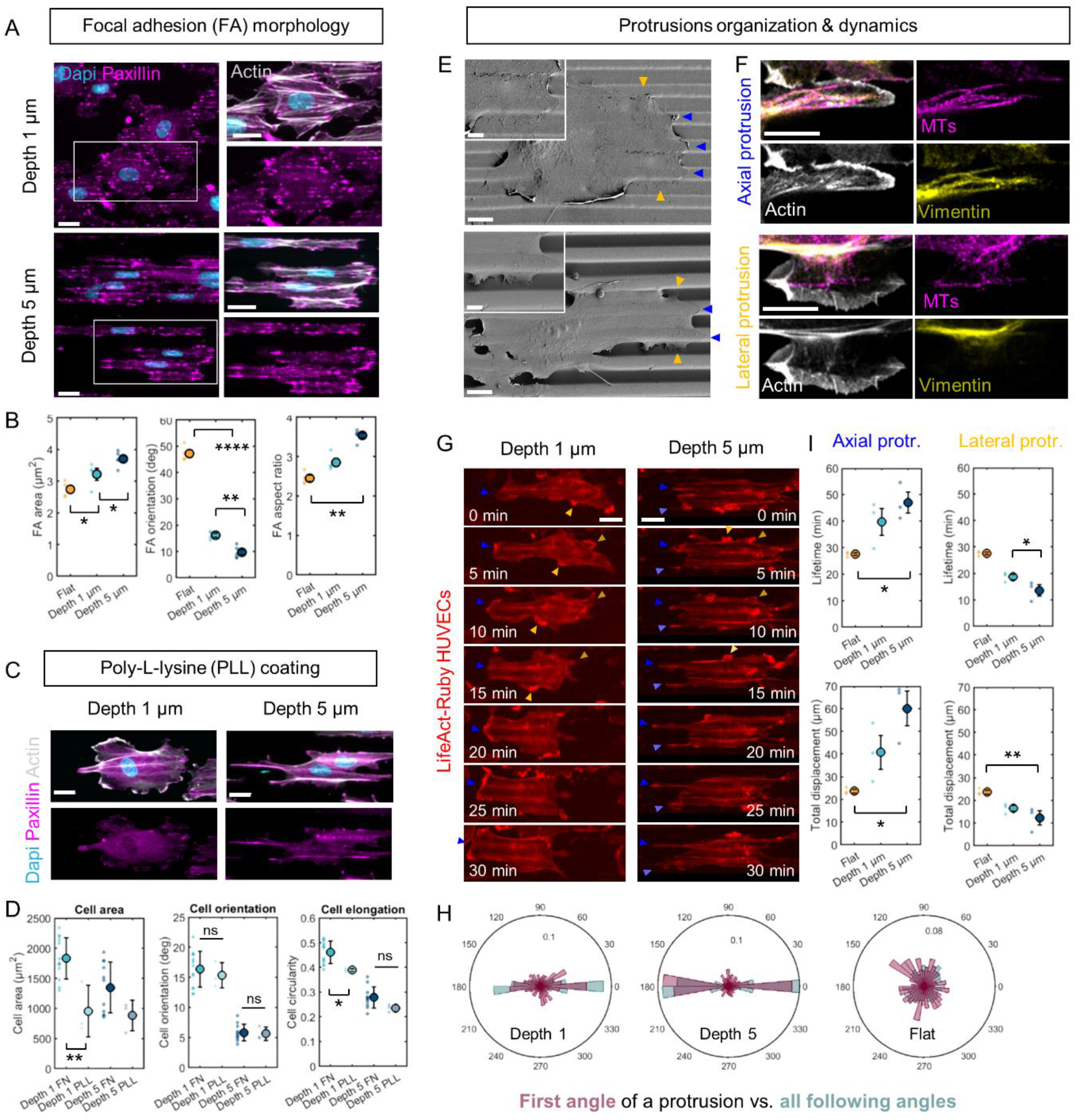
Mechanisms of depth-dependent response to microgrooves in single cells. (A) Immunostaining for paxillin (magenta), actin (grey), and DAPI (cyan) in single cells on 1 μm-deep or 5 μm-deep grooves. Scale bars, 20 μm. Grooves are in the horizontal direction. (B) Quantification of FA area, orientation angle relative to the grooves, and aspect ratio (ratio of major to minor axis). n=4 to 5 independent experiments. (C) Immunostaining for paxillin (magenta), actin (grey), and DAPI (cyan) in single cells on 1 μm-deep or 5 μm-deep grooves coated with poly-L-Lysine (PLL). Scale bar, 20 μm. (D) Quantification of cell area, orientation angle relative to the grooves, and circularity (ratio of minor to major axis). n=3 to 13 independent experiments. (E) Electron microscopy images of single cells on 1 μm-deep or 5 μm-deep grooves. Blue and amber arrowheads represent axial (in the direction of the grooves) and lateral (perpendicular to grooves) protrusions, respectively. Scale bars, 10 μm, 5 μm (insets). (F) Immunostaining for actin (phalloidin, grey), microtubules (β-tubulin, magenta), and intermediate filaments (vimentin, yellow) of axial and lateral protrusions on 5 μm-deep grooves. Scale bar, 10 μm. (G) Images extracted from time-lapse movies of actin-mCherry HUVECs on 1 μm-deep or 5 μm-deep grooves. Scale bar, 20 μm. (H) Quantification of lifetime and total displacement of axial and lateral protrusions. n=3 independent experiments. (I) Polar plots of the distribution of the first angles of protrusions (magenta) or all the following angles of protrusions (blue). For all graphs, dots represent individual experiments. One-way ANOVA, Fisher’s post-test (* p < 0.09, ** p < 0.009 ***, p < 0.0009, **** p < 0.0001). Error bars represent standard error of the mean (SEM).

To test if FA responses alone can explain the groove depth-dependent contact guidance of ECs on microgrooves, we replaced the fibronectin coating with a poly-L-lysine (PLL) coating which allows only weak cell interaction with the substrate. On PLL coating, ECs on microgrooves (especially 5 μm-deep grooves) developed long, filopodia-like protrusions that were oriented in the groove direction (Fig. 6C) but were devoid of FAs and therefore not focally anchored to the substrate. Consistent with this observation, paxillin staining was diffuse with very few punctate FAs (Fig. 6C). In this case, although the overall cell spreading was reduced, cell alignment and elongation on microgrooves were similar to (or even more pronounced than) those on fibronectin coating (Fig. 6D). In addition, the dependence of cell shape on groove depth seen with fibronectin coating persisted in the case of PLL, indicating that FA responses are not necessary to explain the groove depth detection in single ECs. Thus, even without specific FA-mediated interaction with the substrate, ECs are still able to respond to microgrooves, albeit possibly through a different mechanism that may involve a more “passive” or “physical” guidance of filopodia and protrusions.

We next investigated the role of cell protrusions in the initial groove depth-dependent response to fibronectin-coated microgrooves. In electron-microscopy images, membrane protrusions were clearly visible all along the cell borders (Fig. 6E). In the direction of the grooves, “axial” protrusions (Fig. 6E, blue arrowheads) appeared constrained and guided along the ridges while “lateral” protrusions perpendicular to the grooves (Fig. 6E, amber arrowheads) were wider and deformed sharply to “drape over” the ridge edge toward the groove surface. In addition, staining for the different cytoskeletal elements showed that these two types of protrusions were structurally different: while they were both actin-rich by definition, lateral protrusions were less rich in either microtubules or intermediate filaments compared to axial protrusions growing on ridges (Fig. 6F).

We reasoned that these cell protrusions could be at the core of groove depth detection in single cells. More specifically, we hypothesized that the combination of the strong guidance of axial protrusions along the ridges and the high energetic cost associated with the much larger deformations needed for lateral protrusions to reach a neighboring ridge would favor cell axial spreading and would thus explain the more pronounced cell shape anisotropy (i.e. increased cell elongation and alignment) observed on deeper grooves. To test this hypothesis, we used a Lifeact-mCherry HUVEC cell line to analyze the dynamics of cell protrusions on 1 μm- and 5-μm deep grooves (Fig. 6G and Supplementary videos 1 and 2). We observed that while actin-rich protrusions emanated from all sides of the cells, the lateral protrusions either quickly retracted or turned to become axial protrusions in the direction of the grooves (compare the purple and light blue distributions in Fig. 6H). Analyses of the dynamics of the different protrusions showed that axial protrusions on 5 μm-deep grooves had a significantly longer lifetime and total displacement than axial protrusions on either 1 μm-deep grooves or on flat controls (Fig. 6I). Conversely, lateral protrusions on deeper grooves exhibited the shortest lifetime and the smallest displacement. These results confirm that cell protrusions are key elements in the detection of and groove depth-dependent response to substrate topography.

### Cell-cell junctions are not responsible for the decreased contact guidance in endothelial monolayers

We next explored the mechanisms underlying the progressive loss of cell alignment and elongation on microgrooves with increasing cell density. In the case of monolayers, the decreased protrusive activity along with the disassembly of FAs (cf: Fig. 4) point towards a shift in how the cells interact with their substrate. This shift may be attributable to different factors. The first obvious candidate is cell-cell junctions, which mature with increasing time and monolayer density. We initially reasoned that the in-plane tensile and compressive forces mediated by mature cell-cell junctions might compete with and ultimately outweigh cell-substrate adhesion forces, thereby weakening the contact guidance response. To test the influence of cell-cell junctions on the response of EC monolayers to microgrooves, we treated 24 h and 72 h monolayers on 5 μm-deep grooves for 2 h with a cadherin blocking antibody (Huveneers et al., 2012), which completely disassembled cell-cell junctions (Fig. 7A). In low-density monolayers at 24 h of culture, disassembly of cell-cell junctions had little effect on cell shape or alignment (Fig. 7B,C and Supplementary Fig. 5). In 72 h high-density monolayers, however, treatment with the cadherin blocking antibody resulted in a significant loss of cell alignment in the direction of the grooves (Fig. 7B,C). Although the mean cell elongation was not significantly affected, the loss of cell-cell junctions led to a bimodal response where individual cells or groups of cells became alternately either more round or more elongated than the untreated controls.

**Figure 7:**
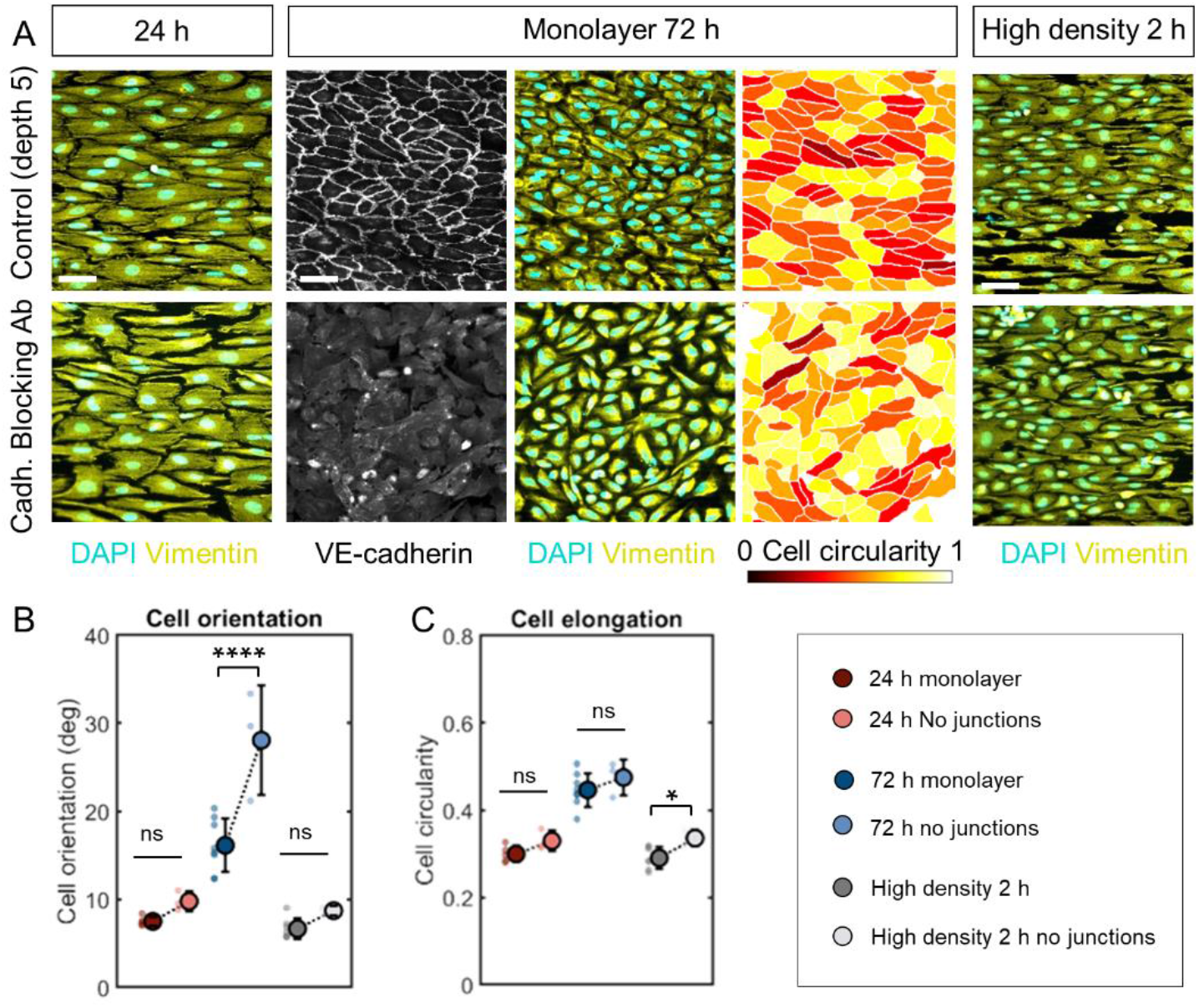
Influence of cell-cell junctions on the response of monolayers to microgrooves. (A) Immunostaining for vimentin (yellow) and DAPI (cyan) or VE-cadherin (grey) for low-density monolayers (24 h of culture), high-density monolayers (72 h of culture), or monolayers after 2 h of culture with or without incubation with a cadherin-blocking antibody on 5 μm-deep grooves. For 72 h monolayers, the right panel represents segmented cells color-coded for cell circularity (1 = round, 0 = elongated). Grooves are in the horizontal direction. Scale bar, 50 μm. (B-C) Quantification of cell orientation angle relative to the grooves and cell circularity (ratio of minor to major axis). Dots represent individual experiments. n=3 to 8 independent experiments. One-way ANOVA, Fisher’s post-test (* p < 0.04, ** p < 0.002, *** p<0.0003, **** p < 0.0001). Error bars represent standard deviations.

To once again separate the effects of culture time and cell density, we also inhibited the formation of cell-cell junctions in monolayers after 2 h of culture (obtained by seeding cells at very high density) (Fig. 7A). In that case, the absence of junctions also resulted in slightly less aligned and less elongated cells on 5 μm-deep microgrooves compared to untreated cultures (Fig. 7B,C and Supplementary Fig. 5). These findings further show that a high cellular density does not prevent cell alignment and elongation, ruling out the possibility that a physical crowding effect is responsible for the loss of contact guidance response to microgrooves with time. Overall, these results suggest that contrary to our hypothesis, the presence of mature cell-cell junctions reinforces rather than weakens the EC contact guidance response on microgrooves. Close inspection of the cell-cell junction staining revealed that junctions perpendicular to the groove direction exhibited a more “zipper-like” morphology as opposed to the more linear aspect of junctions in the groove direction, and this was especially true in the case of moderately confluent monolayers after 24 h of culture (Supplementary Fig. 6). These “zipper-like” or “finger-like” junctions have been shown to be under tension (Hayer et al., 2016; Huveneers et al., 2012), which suggests that the formation of junctions is associated with the establishment within monolayers of anisotropic tensional forces in the direction of the grooves, potentially maintaining and even reinforcing the cell shape anisotropy initially present in single cells.

### Cell density-dependent loss of response to microgrooves is principally determined by *de novo* basement membrane secretion

The density and composition of the ECM on which cells are cultured *in vitro* have a strong influence on cell shape (Mooney et al., 1992; Reinhart-King et al., 2005). We wondered if the loss of contact guidance response to topography in EC monolayers was related to *de novo* secretion of BM proteins by the cells. Immunofluorescence staining demonstrated that with increasing time and cell density, the fibronectin coating used in the preparation of our substrates was progressively replaced by proteins secreted by the cells themselves (Fig. 8A). More specifically, fibronectin, collagen IV, and laminin staining at different times of culture showed that a dense matrix progressively accumulated on the surface of the substrate, even to the point of partially filling the grooves in 72 h cultures as shown in cross-sections obtained by confocal microscopy (Fig. 8A,B). This BM secretion is determined principally by time in culture rather than cell density (Supplementary Fig. 7). We hypothesized that this secreted BM may impair the interaction of cells with the microgrooves, explaining the loss of contact guidance response. To test this hypothesis, we first detached 72 h monolayers from the substrate without removing the secreted BM and seeded new cells at low density on this matrix (Fig. 8C). Single cells cultured for only 2 h on the 72 h-secreted matrix exhibited an alignment and elongation similar to cells in 72 h monolayers, significantly lower than in the control 2 h condition (Fig. 8D,E), showing that cell elongation and alignment are determined by the underlying secreted matrix rather than by the microgroove architecture.

**Figure 8:**
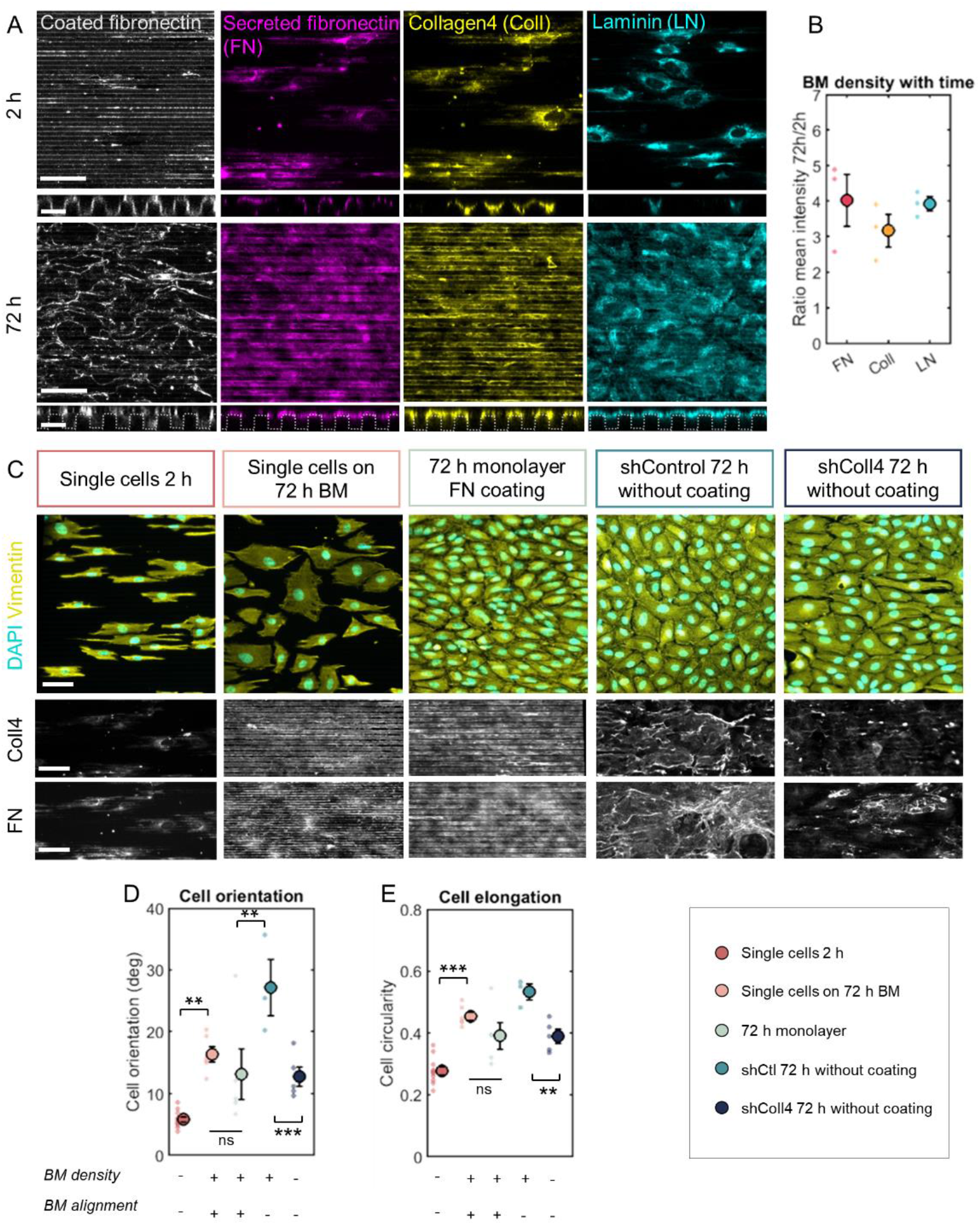
Influence of extracellular matrix secretion on the response of monolayers to microgrooves. (A) Coated fibronectin visualized by fluorescent fibrinogen and immunostaining against secreted fibronectin (magenta), collagen IV (yellow), and laminin (cyan). Bottom panels show Z cross-sections. Scale bars, 50 μm, 10 μm (cross-sections). (B) Ratio of the mean fluorescence intensity at 72 h to that at 2 h for the different ECM proteins. (C) Immunostaining for vimentin (yellow) and DAPI (cyan) and the corresponding secreted basement membrane stained for collagen IV or fibronectin (grey) for single cells (2 h of culture), single cells cultured on a basement membrane secreted for 72 h, and high-density monolayer (72 h of culture), shControl-or shCollagenIV-transfected cells cultured for 72 h on uncoated microgrooves. Grooves are in the horizontal direction. Scale bar, 50 μm. (D-E) Quantification of cell orientation angle relative to the grooves and cell circularity (ratio of minor to major axis). Dots represent individual experiments. n=3 to 11 independent experiments. One-way ANOVA, Fisher’s post-test (* p < 0.02; ** p < 0.009; *** p<0.0009 ****, p < 0.0001). Error bars represent standard error of the mean (SEM).

To further explore the role of the secreted BM, we transfected HUVECs with an shRNA against collagen IV α1 (shColl4) which impairs the secretion and assembly of BM (Bignon et al., 2011). As coating with fibronectin is known to rescue defects in deposition of mutated collagen IV and other matrix components (Ngandu Mpoyi et al., 2020; Umana-Diaz et al., 2020), we cultured these cells (as well as the shRNA control cells) on uncoated microgrooved substrates. While the secreted BM on fibronectin-coated substrates tended to follow the orientation of the grooves, the secreted BM of shControl cells on uncoated substrates was dense but adopted a random orientation (Fig. 8C). Interestingly, shControl cells on uncoated substrates after 72 h of culture were significantly less aligned and elongated than 72 h monolayers on fibronectin-coated microgrooves (Fig. 8D,E), confirming that cells respond principally to the secreted BM. In shColl4 cultures, the secreted collagen IV and fibronectin were significantly less dense after 72 h of culture than those in the shControl cultures (Fig. 8C). As expected in that case, shColl4 cells exhibited more pronounced cell elongation and alignment than the shControl cells (Fig. 8D,E).

Taken together, these findings demonstrate that the secreted BM is a crucial determinant of EC response to topography in high cell density cultures. While cells at low density or short culture time respond to the underlying substrate topography, they progressively switch to interact principally with their own secreted matrix as cell density and time in culture increase. If this matrix is sufficiently dense and exhibits an orientation that is different from that of the underlying substrate, cells lose the response to substrate topography.

## Discussion

In this study, we investigated the contact guidance response of vascular ECs to microgroove substrates not only in single cells at early times of culture as was done in most previous studies but also later in time as cells attain confluence, form monolayers, and exhibit collective behavior. The response to substrate topography of monolayers compared to single cells has, to our knowledge, not been previously addressed but is crucial as vascular ECs in blood vessels are naturally found as monolayers. Monolayer response to microgrooves is also important in the context of endothelialization of implantable endovascular devices whose surfaces may be patterned in order to optimize performance.

We used substrates composed of parallel arrays of grooves in the micrometric range (2-5 μm in width and depth). These structures constitute idealized mimics of the anisotropy often found in native BMs. This anisotropy can be found not only in the fibrillar orientation of BM components at the nanometric scale but also in the micrometric topography that takes the form of undulations due to BM supramolecular aggregates (Leclech et al., 2020). While contact guidance responses have been observed at both scales, the underlying mechanisms and the subcellular structures involved may be different for the two scales. Precise characterization of the topographical organization of BMs *in vivo* remains largely missing and will be key for engineering more physiologically relevant substrates *in vitro*.

The cellular and subcellular mechanisms underlying contact guidance remain unclear, and different subcellular structures have been proposed in the literature as key players in this process (Leclech and Barakat, 2021). Most of the models proposed thus far centrally implicate FAs and the actin cytoskeleton. In our system, individual HUVECs cultured at short time on microgrooves exhibited FAs that are aligned and elongated in the direction of the grooves, as described in other studies (Franco et al., 2011; Ray et al., 2017; Saito et al., 2014). In addition, FAs were more aligned and elongated on deeper (5 μm) grooves where the most pronounced contact guidance was observed. The principal mechanism evoked in the literature in the case of single cells considers lateral constraints on FA growth as the main trigger of contact guidance (Ray et al., 2017), especially on ridge widths that are comparable to the lateral dimensions of mature FAs (sub-micron to a couple of microns). These lateral constraints may be expected to be especially pronounced at the ridge edges where most FAs localize, in agreement with the “discontinuity” theory of Curtis and Clark who proposed that sharp discontinuities in the substrate (e.g., edges of grooves) can induce FA formation and actin condensation (Curtis and Clark, 1990). However, in this scenario, the dependence of FA morphology on groove depth remains unclear and could therefore be a consequence of rather than a cause for cell elongation. To test the central role of FAs, we used a poly-L-lysine surface coating that prevents the formation of mature FAs. Single ECs on this substrate formed long protrusions and retained a groove depth-dependent alignment and elongation similar to that observed in the presence of FAs. These results indicate that contact guidance on microgrooves can occur even without FAs, albeit possibly through other mechanisms whose physiological relevance remains to be established. It also suggests the possible existence of a “passive”, non-adhesive component to the cell alignment and/or elongation associated with contact guidance, although the precise nature of such an interaction remains to be elucidated.

The role of cell protrusions, while more debated in the literature, has also been put forth to explain cell response to substrate anisotropy (Franco et al., 2011; Fujita et al., 2009). In our system, the clear delineation between long and stable protrusions in the groove direction and short and highly transient protrusions perpendicular to the groove direction suggests that cell protrusions are key drivers of initial cell alignment and elongation. In accordance with this notion, the increased deformation of protrusions that would occur in the case of deeper grooves would further favor axial protrusions over lateral protrusions, potentially explaining the observed increase in EC elongation and alignment for deeper grooves. In support of this idea, membrane deformation has been proposed as a mechanism involved in the detection and response to high aspect-ratio structures (Leclech and Villard, 2020; Lou et al., 2018). Although these findings shift our thinking about contact guidance in single cells from a FA-centered to a protrusion-centered model, it should be noted that the roles of these two structures are likely intricately intertwined. More specifically, they probably act synergistically with cell protrusions guiding FA growth and FAs anchoring and stabilizing cell protrusions. Better understanding of the precise dynamics of these two elements and their interactions is needed to determine the exact sequence of events involved in the early detection of substrate topography.

Anchored to FAs, the actin cytoskeleton is widely considered to be a central player in contact guidance responses by generating the anisotropic intracellular forces leading to overall cell alignment and elongation (Leclech and Barakat, 2021; Ray et al., 2017). In our system, the actin cytoskeleton in single ECs exhibited a striking organization in the form of thick stress fiber bundles localized above the grooves and bridging the FAs present on the sides of adjacent ridges. While microtubules and intermediate filaments primarily followed the cell orientation, actin conformed more closely to the grooves, suggesting a role in the detection and response to topography. Thus, the observation that the contact guidance response in single ECs remains the same upon either inhibition of actomyosin-mediated contractility with blebbistatin or actin depolymerization with latrunculin A and cytochalasin D came as a surprise. These results raise questions about the central role of actin in the morphological changes associated with contact guidance in single cells. Other studies have reported similar results and proposed that the initial contact guidance response is actin- and contractility-independent (Franco et al., 2011; Sales et al., 2017). Our findings suggest, however, that actin may play a more important role in monolayers, possibly through its interaction with cell-cell junctions (Hoelzle and Svitkina, 2012).

The role of other cytoskeletal elements in contact guidance has been largely overlooked thus far. We demonstrated here an important role for microtubules in controlling cell elongation, which may be linked to the mechanical rigidity of this network. Disruption of contact guidance upon nocodazole treatment has also been reported in other cell types (Lee et al., 2016; Tabdanov et al., 2018) but not specifically the alteration of cell elongation. We have not addressed here the role of intermediate filaments, which may potentially be important as well. Overall, disruption of the different cytoskeletal elements never completely abolished the contact guidance response of ECs. This may be due to incomplete cytoskeletal disruption, compensatory mechanisms by the other cytoskeletal elements, or indicative of a more passive component of the response that is independent of the cytoskeleton or other intracellular elements. It is also interesting to point out that pharmacological disruption of the different cytoskeletal elements (as well as the increase in cell density, cf Fig. 2) has a more pronounced effect on cell elongation than on cell alignment. It is possible that these two morphological responses are actually driven by different intracellular elements and mechanisms. In this context, cell alignment appears to be the most conserved feature and may therefore constitute the “passive” portion of contact guidance, while cell elongation is a more active process that relies upon the cytoskeleton (microtubules in particular). In support of this hypothesis, overall cell alignment has been reported to occur as early as 10 minutes after seeding, well before cell elongation or the alignment of intracellular structures (Sales et al., 2017).

The main novelty of this study was to follow the contact guidance response of ECs in time, from single cells to the formation and maturation of monolayers. We characterized the changes in intracellular organization associated with this transition and observed a disassembly of central FAs accompanied by a relocalization of actin stress fibers to the cell periphery. These changes are very likely driven by the formation of cell-cell junctions, which are known to be physically associated with the actin cytoskeleton (Hoelzle and Svitkina, 2012; Millán et al., 2010) and with FA-associated proteins (Huveneers et al., 2012). By tracking EC shape on microgrooves in time and with increasing cell density, we found that cell alignment and elongation progressively decrease, which we interpret as a gradual loss of response to substrate topography. The disassembly of FAs in monolayers supports the idea of a weakened, or at least altered, interaction of cells with the substrate. We initially thought that in light of the commonly evoked competition between cell-substrate interaction and cell-cell interaction in monolayers (Maruthamuthu et al., 2011; Noethel et al., 2018; Zimmermann et al., 2016), the presence of cell-cell junctions would change the balance of forces, increasing “planar” tension and decreasing the interaction with the basal substrate. Supporting this idea, disassembly of FAs has been observed in epithelial cells under strain (Noethel et al., 2018). However, what we observed is that disassembly of cell-cell junctions with a cadherin blocking antibody did not restore cell alignment and elongation in monolayers but on the contrary tended to further decrease the contact guidance response. This result indicates a more complex role for cell-cell junctions than expected. We hypothesize that cell-cell junctions, when formed between cells that are already elongated, actually provide an anisotropic tensional force that reinforces contact guidance. This notion is supported by the observation that in low-density monolayers, the junctions along the major axis of the cell (parallel to the grooves) are more linear compared to more “zig-zag” junctions along the minor axis of the cell (perpendicular to grooves). In high-density monolayers, the high variability of cell elongation patterns observed upon disassembly of junctions possibly reflects the complexity of the pattern of forces present within the monolayer, with the coexistence of tensional and compressive forces.

Another consideration in elucidating the interactions of cell monolayers with substrate topography is the *de novo* secretion by the cells of ECM proteins. While this process has thus far been largely overlooked in contact guidance studies, we demonstrate here that it is a crucial determinant of cell morphology in monolayers. Indeed, the progressive deposition of a dense matrix by cells with time and increasing cell density masks the underlying substrate topography, thereby decreasing contact guidance. Our results show that EC shape is tightly controlled by the underlying deposited matrix: when it is dense and/or randomly oriented, cells are more round and less aligned than in regions where the matrix is less dense and follows the orientation of the grooves.

In addition to shedding valuable insight into the mechanisms of contact guidance in both single cells and monolayers, the results of the present study may have important implications for the endothelialization of implantable endovascular devices such as stents and grafts. A promising strategy for improving endothelialization is microstructuring implant surfaces (typically with grooves) to guide cell migration and control proliferation (Sprague et al., 2012; Tan et al., 2017; Vesga et al., 2017; Wang et al., 2019). In this context, the present results suggest that individual cells that initially colonize the implant will strongly respond to the surface topography. Upon cell proliferation and monolayer formation, cells will progressively secrete their own BM and begin to respond to it, thereby attenuating the interaction with the implant surface. A potentially positive consequence of the gradual loss of contact guidance is that the cells may then become more responsive to the native extracellular cues such as flow, BM topography, and substrate stiffness.

As a general conclusion, we provide here a vivid illustration of the strong influence of substrate topography on the regulation of EC shape and alignment. A key contribution of the present work is the documentation of the evolution of the contact guidance response as single ECs proliferate to ultimately form highly confluent monolayers. The identification of different regimes and distinct mechanisms of response to substrate topography depending on cell density reveal fundamental differences between single cells and monolayers that will undoubtedly lead to more careful examination of the specificity of cellular monolayers in future contact guidance studies.

## Conclusions

In this study, we have highlighted different regimes and mechanisms of response to substrate topography depending on time and cell density. Single cells interact strongly with their underlying substrate, and are able to detect microstructures and their dimensions thanks principally to cell protrusions. Our results challenge the commonly evoked central role of FAs and the actin cytoskeleton in the initial steps of contact guidance. Alternatively, we describe an important role for cell protrusion guidance and the microtubule network in controlling cell shape in response to microgrooves. During monolayer formation and maturation, we found that cells progressively switch from interacting with the microgrooves to interacting with their own secreted BM, thereby losing the contact guidance response. This transition is associated with changes in the organization of FAs and the actin cytoskeleton, highlighting a different mode of interaction with the substrate and regulation of cell shape within monolayers.

## Materials & Methods

### Fabrication of microgrooved substrates

The original microstructured silicon wafer was fabricated using classical photolithography procedures by UV illumination (MJB4 Mask Aligner, 23 mW/cm^2^ power lamp, SUSS MicroTec, Germany) of a layer of SU8-2010 (MicroChem, USA) through a hard chromium mask. After exposure to trichloro(1H,1H,2H,2H-perfluorooctyl)silane (Sigma) vapor for 20 min, the silicon wafer was used to create polydimethylsiloxane (PDMS Sylgard 184, Sigma Aldrich, ratio 1:10) replicates. To create the final coverslip on which the cells were cultured, liquid PDMS was spin coated at 1500 rpm for 30 s on the PDMS mold. Before reticulation overnight at 70°C, a glass coverslip was placed on top of the PDMS layer. After reticulation, the glass coverslip attached to the microstructured PDMS layer was gently demolded with a scalpel and isopropanol to facilitate detachment. Microstructured coverslips were then sonicated for 10 min in ethanol for cleaning and finally rinsed with water.

### Cell culture

Prior to cell seeding and after a 30 s plasma treatment, the microgroove substrates were incubated for 1 h with 50 μg/ml fibronectin solution (Sigma) at room temperature. For visualization of the fibronectin localization, the fibronectin was mixed with fluorescent fibrinogen 647 (Thermofisher F35200). Poly-L-lysine (Sigma P4707) coating was prepared similarly by incubating the substrates with a 0.003% solution diluted in water for 1 h at 37°C. Coverslips were then washed 3 times with PBS before culture. In conditions without coating, cells were directly seeded on the coverslips after the plasma treatment. Human umbilical vein endothelial cells (HUVECs, Lonza) in passages 4-8 were cultured in EGM2-MV medium (Lonza) at 37°C in a humidified atmosphere of 95% air and 5% CO_2_. At confluence, cells were detached with trypsin (Gibco, Thermo Fisher Scientific) and seeded onto either control PDMS substrates or microgroove coverslips at densities of 30,000-50,000 cells/cm^2^.

To remove the cells while maintaining the secreted BM, 72 h monolayers were treated for 30 min to 1 h with 20 mM ethylenediaminetetraacetic acid (EDTA, Sigma) to detach the cells. The cultures were then gently washed with PBS to remove the detached cells, and new cells were directly seeded on the surface of the coverslips. The presence of an intact BM at the surface of the coverslip was verified by immunostaining for fibronectin and collagen IV.

### Transfected HUVEC cell lines

ECs isolated from a single umbilical cord were split in two and transduced before the first passage with lentivirus encoding control or COL4A1 targeting shRNA (Sigma-Mission shRNA) (Bignon et al., 2011). Transduced cells were selected using puromycin and amplified before storage in liquid nitrogen. Experiments using thawed cells were performed until passage 5. ECs expressing LifeAct-mRuby were generated as described elsewhere (Atlas et al., 2021).

### Cytoskeletal and cell-junction pharmacological treatment

The following cytoskeletal pharmacological agents were used: blebbistatin (Sigma B0560) at 100 μM, latrunculin A (Millipore 428026) at 10 nM, cytochalasin D (Sigma C2618) at 20 nM, and nocodazole (Sigma M1404) at 0.2 μM. The incubation times were 90 min for all pharmacological agents. Controls were treated with the equivalent DMSO concentration. Inhibition of cell-cell junctions was achieved using a VE-cadherin function-blocking antibody (clone 75, BD Biosciences 610251) in which the cells were incubated at 12.5 μg/ml for 2 h before fixation.

### Immunostaining

After different times of culture (2 h, 24 h, 48 h or 72 h), culture coverslips were fixed with 4% paraformaldehyde (Thermo Fisher) in PBS for 15 min. After 1 h in a blocking solution containing 0.25% Triton and 2% bovine serum albumin (BSA), cultures were incubated for 1 h at room temperature with primary antibodies as follows: rabbit anti-VE-cadherin (ab33168, Abcam), mouse anti-paxillin (MA5-13356, Thermofisher), mouse anti-vimentin (ab8069, Abcam) or rabbit anti-vimentin (ab92547, Abcam), mouse anti α-tubulin (Sigma, T5168), mouse anti-fibronectin (ab6328, Abcam), rabbit anti-collagen IV (ab6586, Abcam), or rabbit anti-laminin (ab11575, Abcam). All antibodies were diluted 1/400 - 1/200 in a solution containing 0.25% Triton and 1% BSA. Coverslips were washed three times with PBS and incubated for 1 h at room temperature with Alexa Fluor 555-conjugated donkey anti-rabbit antibody (ab150074, Abcam) or Alexa Fluor 488-conjugated donkey anti-mouse antibody (ab150105, Abcam) and DAPI. When needed, actin was stained in this last step using phalloidin (LifeTechnologies).

### Microscopy

#### Epifluorescence and confocal microscopy of fixed samples

Epifluorescence images were acquired on an inverted microscope (Nikon Eclipse Ti) with a 20X objective (Nikon Plan Fluor NA=0.5). Confocal microscopy images were acquired on an inverted TCS SP8 confocal microscope (Leica) using a 63X objective.

#### Scanning electron microscopy

Cultures were fixed in 2% glutaraldehyde in 0.1 M phosphate buffer at pH 7.4. They were dehydrated in a graded series of ethanol solutions and then dried by the CO_2_ critical-point method using EM CPD300 (Leica Microsystems). Samples were mounted on an aluminum stub with a silver lacquer and sputtercoated with a 5 nm platinum layer using EM ACE600 (Leica Microsystems). Acquisitions were performed using a GeminiSEM 500 (Zeiss).

#### Live cell imaging

Live recordings of Lifeact-mCherry HUVECs were performed after 2 h of culture using an automated inverted microscope (Nikon Eclipse Ti) equipped with temperature and CO_2_ regulation and controlled by the NIS software (Nikon). Images were acquired with a 20X objective (Nikon Plan Fluor NA=0.5) for 3 h at 5 min intervals. Tracking of the different cell protrusions was performed manually using the MTrackJ plugin in Fiji. Protrusions were defined as actin-rich structures growing from the main cell body and were tracked from the moment they emerged from the cell outline until they completely retracted, were joined by the cell body moving forward, or changed direction to become axial protrusions in the case of lateral protrusions.

### Data analysis

#### Automated cell shape analysis

Cell shape in fixed monolayers was automatically analyzed by first segmenting the cell outline using either the “Tissue Analyzer” plugin in Fiji or the CellPose algorithm in Python (Stringer et al., 2021). Different cell shape parameters and associated color-coded visual representations were subsequently generated using a custom-made Matlab code. Cell orientation was defined as the mean absolute value of the angle between the major axis of the ellipse that best fits the cell contour and the microgroove direction; 0° corresponds to the microgroove direction and 90° is orthogonal to the microgrooves. Cell circularity was defined as the ratio of the minor-to-major axis of the best fit ellipse of the cell.

#### Analysis of cytoskeletal organization

Cytoskeletal orientation was analyzed using the “OrientationJ” plugin in Fiji. Normalization was done by the sum of all values. The localization of the different cytoskeletal elements within a cell was analyzed using a custom-made Fiji plugin. Briefly, a line perpendicular to the nucleus major axis and passing through the nucleus center was drawn across the cell, and the fluorescence intensity profile of the cytoskeletal elements was then plotted along this line and normalized by the sum of the values.

#### Focal adhesion analysis

FA morphometric analysis was performed using a custom-made Fiji macro similar to the one described by Natale et al., 2019. Briefly, a blurred image of the FAs was subtracted from the original images. The images were further processed using thresholding to obtain binary images. Pixel noise was eliminated using the ‘erode’ command, and particle analysis was then implemented to extract the morphometric descriptors.

### Statistical analysis

All analyses are based on at least 3 independent experiments. Statistical analyses were performed using the GraphPad Prism software. Multiple groups with a normal distribution were compared by a one-way ANOVA followed by Tukey’s or Fisher’s posthoc test. The number of data points for each experiment, the specific statistical tests, and the significance levels are noted in the corresponding figure legends.

## Author Contributions

C.L. Conceptualization; Methodology; Software; Formal Analysis; Investigation; Writing - Original Draft; Visualization; Funding acquisition. A.K. Investigation. L.M. Conceptualization; Resources. A.I.B. Conceptualization; Writing - Original Draft; Supervision; Project administration; Funding acquisition.

## Acknowledgments & Funding

We thank P. Mahou and the Polytechnique Bioimaging Facility for assistance in imaging. The Polytechnique Bioimaging Facility is supported by Région Ile-de-France (interDIM) and Agence Nationale de la Recherche (ANR-11-EQPX-0029 Morphoscope2, ANR-10-INBS-04 France BioImaging). This work was supported in part by an endowment in Cardiovascular Bioengineering from the AXA Research Fund (to AIB) and postdoctoral fellowships from the Lefoulon-Delalande Foundation and the Bettencourt-Schueller Foundation (to CL).

## Data Availability

The data that support the findings of this study are available from the corresponding author upon reasonable request.

## Supplementary Figures

**Supplementary Figure 1:**
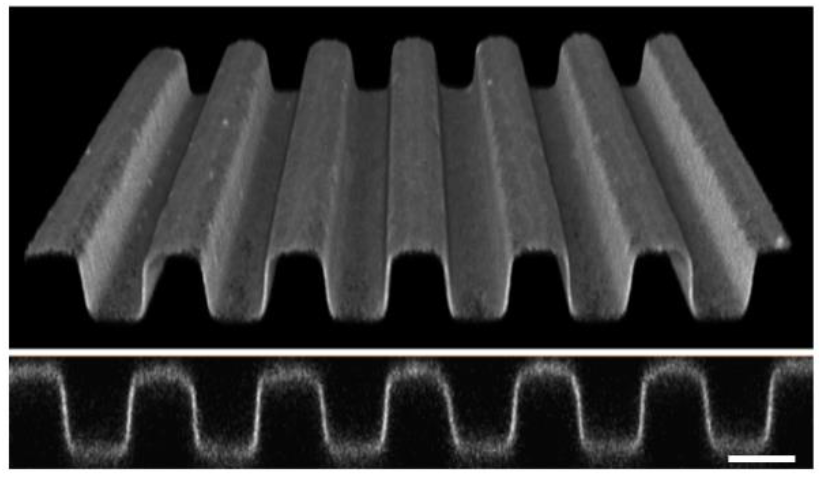
Fibronectin coating on microgrooves. Confocal 3D reconstruction and cross-section of fluorescent fibrinogen showing homogeneity of fibronectin coating on 5 x 5 x 5 μm (w x s x d) grooves. Scale bar 5 μm.

**Supplementary Figure 2:**
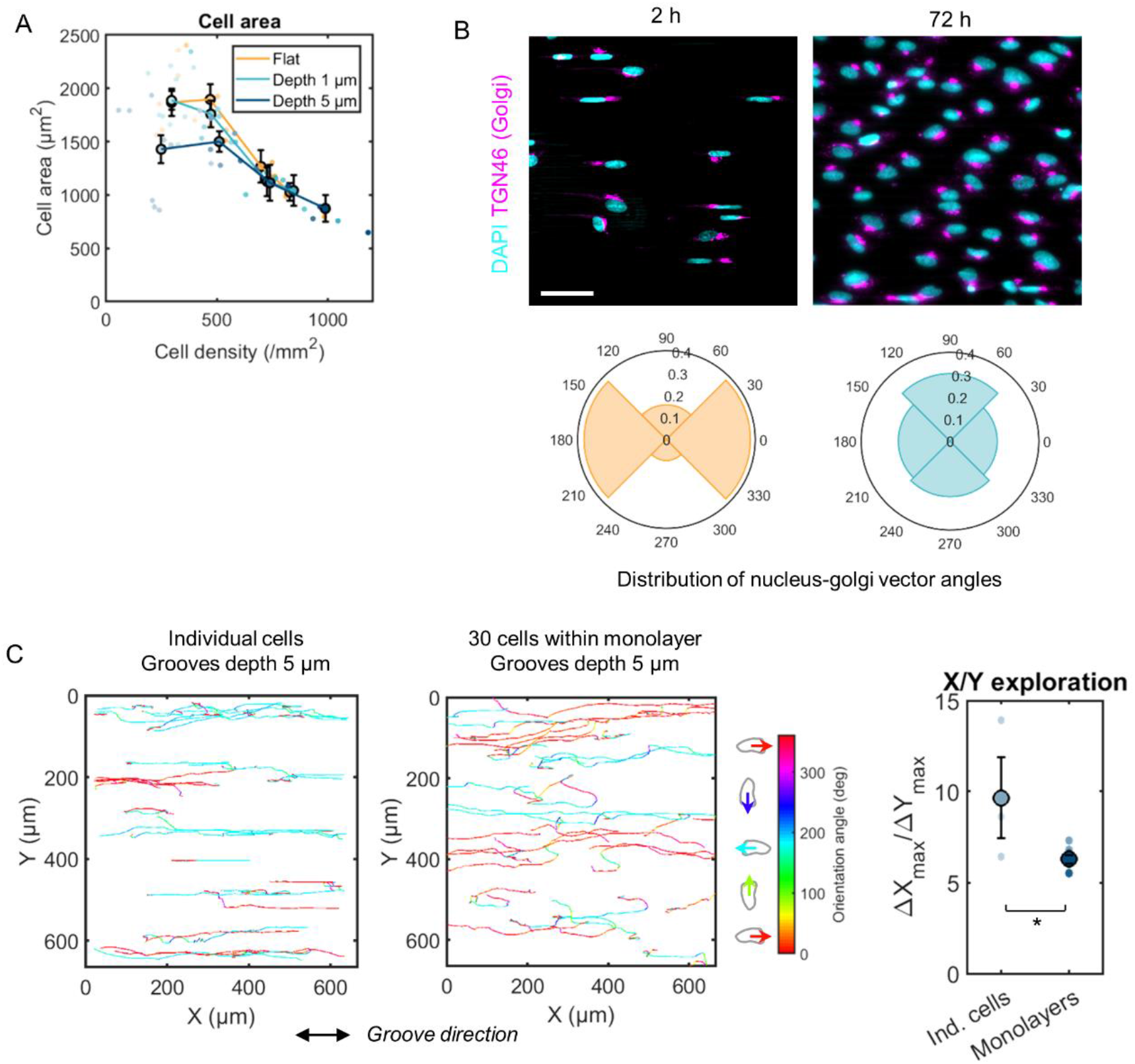
Functional ramifications of the loss of response to microgrooves in monolayers. (A) Quantification of cell area for increasing time and cell density on flat surfaces (yellow), 1 μm-deep grooves (light blue), and 5 μm-deep grooves (dark blue). Dots represent individual experiments, regrouped by density groups corresponding to 2 h, 24 h, 48 h, and 72 h cultures, with at least 3 independent experiments per group. Error bars represent standard error of the mean (SEM). (B) Immunostaining for TGN46 (Golgi, purple) and DAPI (blue) of HUVECs after 2 h (single cells) or 72 h (monolayers). Scale bar, 50 μm. Distribution of the orientation of nucleus-golgi vectors relative to the groove direction (0°). (C) Individual cells (left) or cells in monolayers (right) were recorded for 24 h, with a time interval of 15 min. Trajectories were obtained by tracking the cell nucleus visible with Hoescht. Graphs show the accumulated cell trajectories during the 24 h of recording, color-coded for the orientation angle of each displacement vector. Quantification shows the ratio between x (along grooves) and y (perpendicular to grooves) displacement (calculated as |Xmax — Xmin|/|Ymax — Ymin| for each trajectory). Dots represent individual experiments. n=3 to 8 independent experiments, unpaired t-test (* p < 0.03). Error bars represent standard error of the mean (SEM).

**Supplementary Figure 3:**
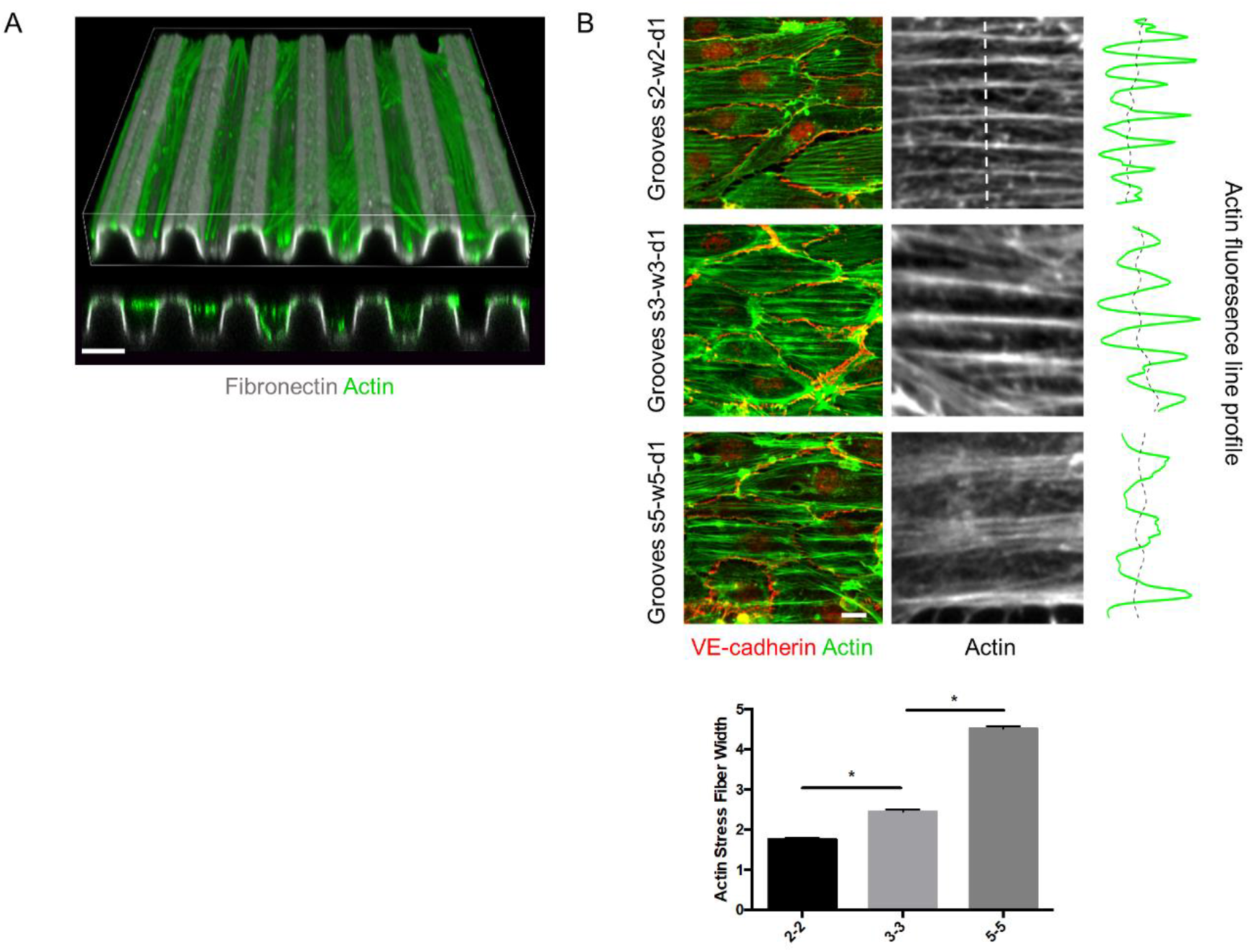
Organization of the actin cytoskeleton relative to the grooves. (A) Confocal 3D reconstruction and cross-section of fluorescent fibrinogen (grey) and actin (green) on 5 x 5 x 5 μm (w x s x d) grooves showing the spatial localization of actin filaments within the grooves. Scale bar 5 μm. (B) Organization of actin stress fibers on grooves of different widths and spacings (2, 3, and 5 μm). Quantification of the width of actin bundles shows a correlation between groove width and actin bundle width which illustrates that the actin network is particularly responsive to the substrate topography.

**Figure supplementary 4:**
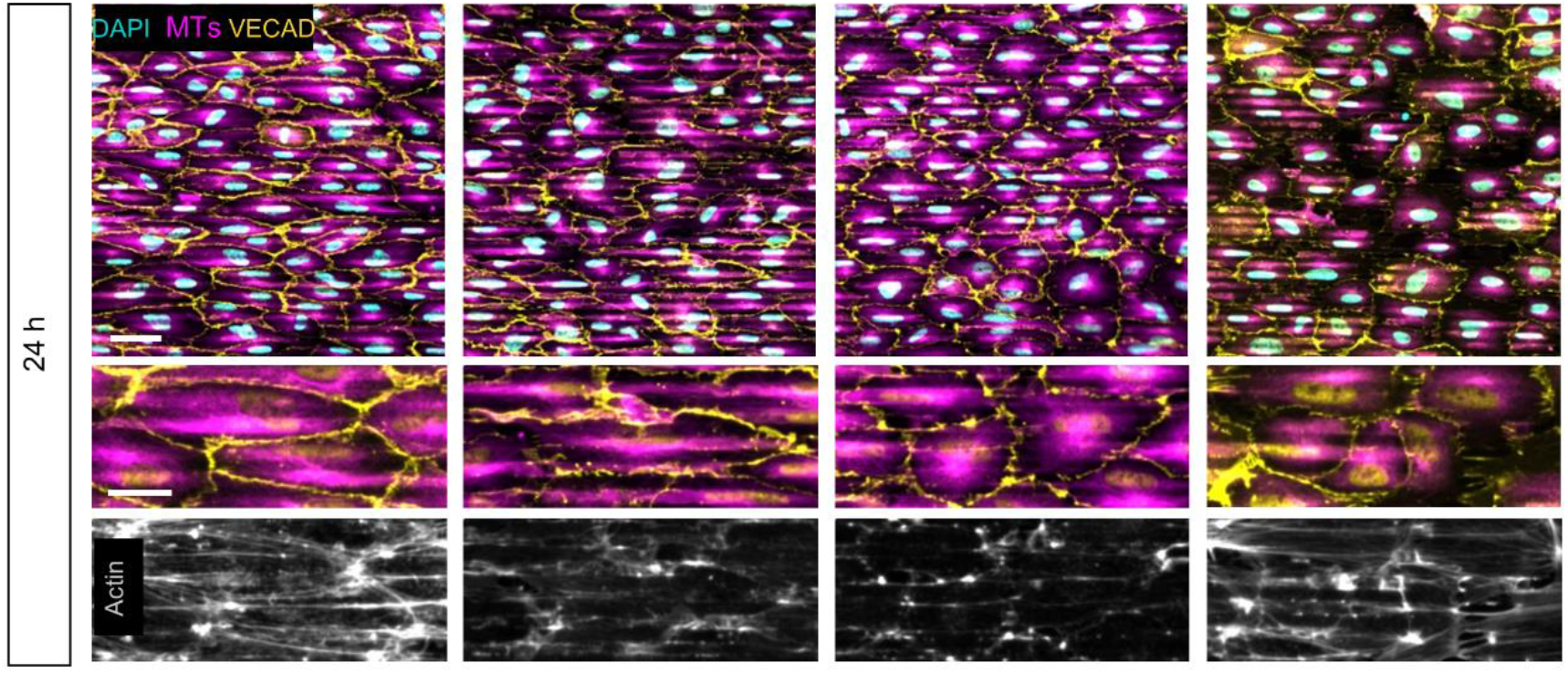
Influence of the different cytoskeletal elements on the response to microgrooves. Immunostaining after 24 h of culture for microtubules (β-tubulin, magenta), actin (phalloidin, gray), DAPI (cyan) and VE-cadherin (yellow) of control cells (+DMSO) or cells treated for 90 min with 100 μM blebbistatin, 0.1 nM latrunculin A + 0.2 nM cytochalasin D or 0.2 μM nocodazole. Scale bars, 50 μm, 25 μm (insets). Grooves are in the horizontal direction.

**Supplementary Figure 5:**
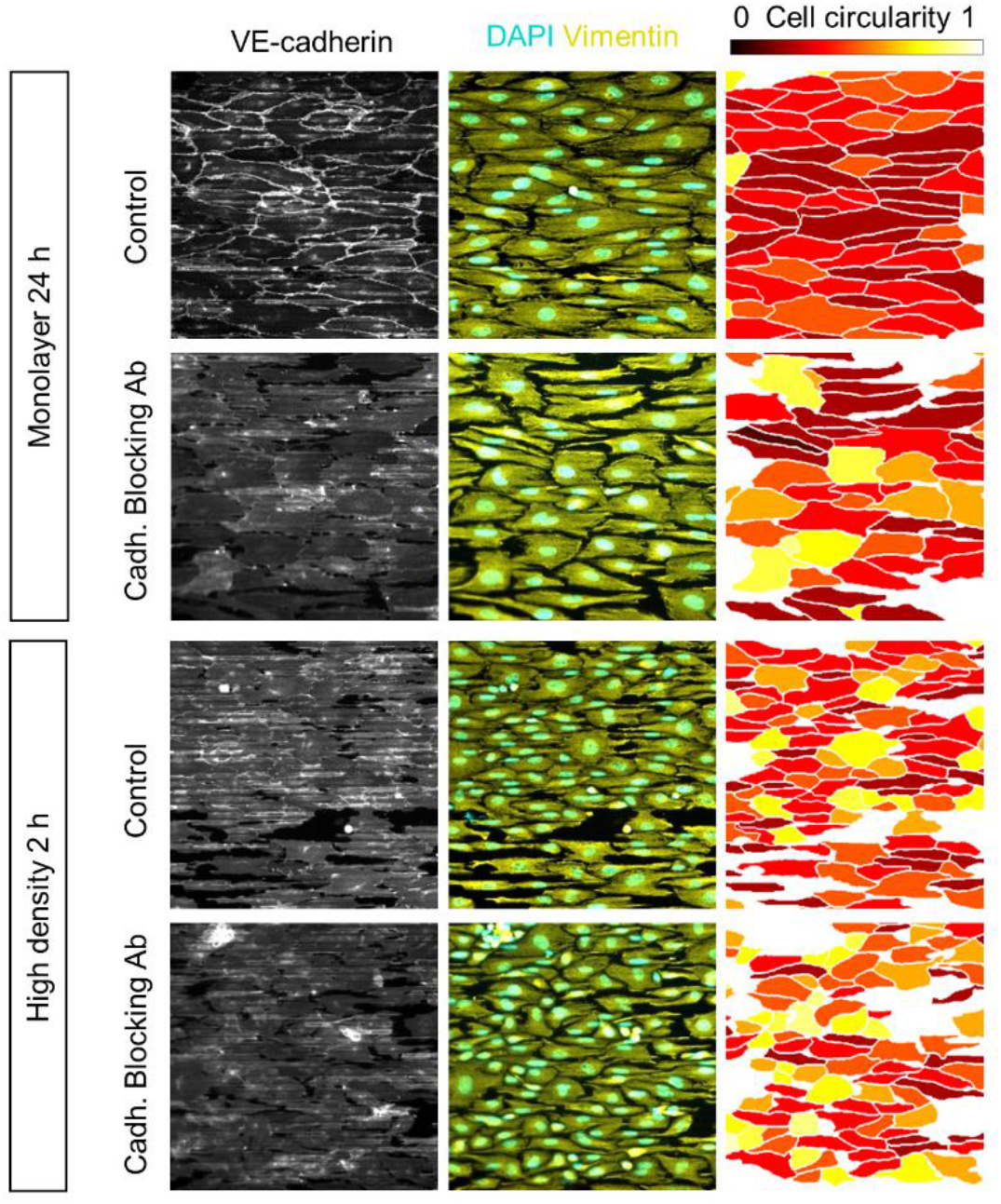
Influence of cell-cell junctions on the response of monolayers to microgrooves. Immunostaining for vimentin (yellow) and DAPI (cyan) or VE-cadherin (grey) for low-density monolayers (24 h of culture) or monolayers after 2 h of culture with or without incubation with a cadherin-blocking antibody on 5 μm-deep grooves. Right panel: segmented cells color-coded for cell circularity (1 = round, 0 = elongated). Grooves are in the horizontal direction. Scale bar, 50 μm.

**Supplementary Figure 6:**
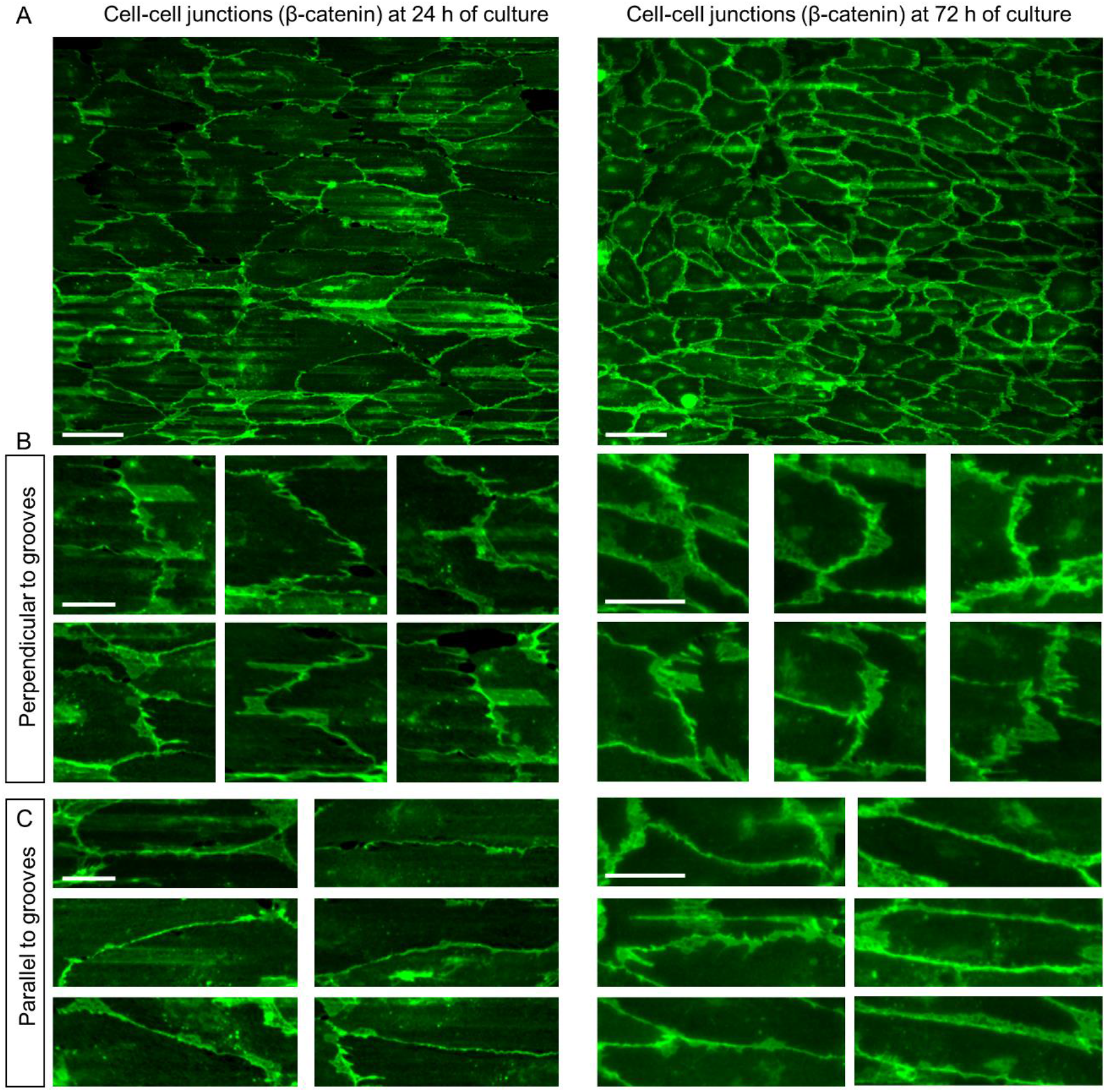
Morphology of cell-cell junctions. (A) HUVEC monolayer after 24 h (left) or 72 h (right) of culture on 5 x 5 x 5 μm (w x s x d) grooves immunostained for β-catenin (green). Scale bar, 50 μm. (B) Zoom-in on cell-cell junctions perpendicular to the groove direction, showing “zipper-like” or “finger-like” junctions. Scale bar, 20 μm (C) Zoom-in on cell-cell junctions parallel to the groove direction, showing more linear junctions. Scale bar, 20 μm.

**Supplementary Figure 7:**
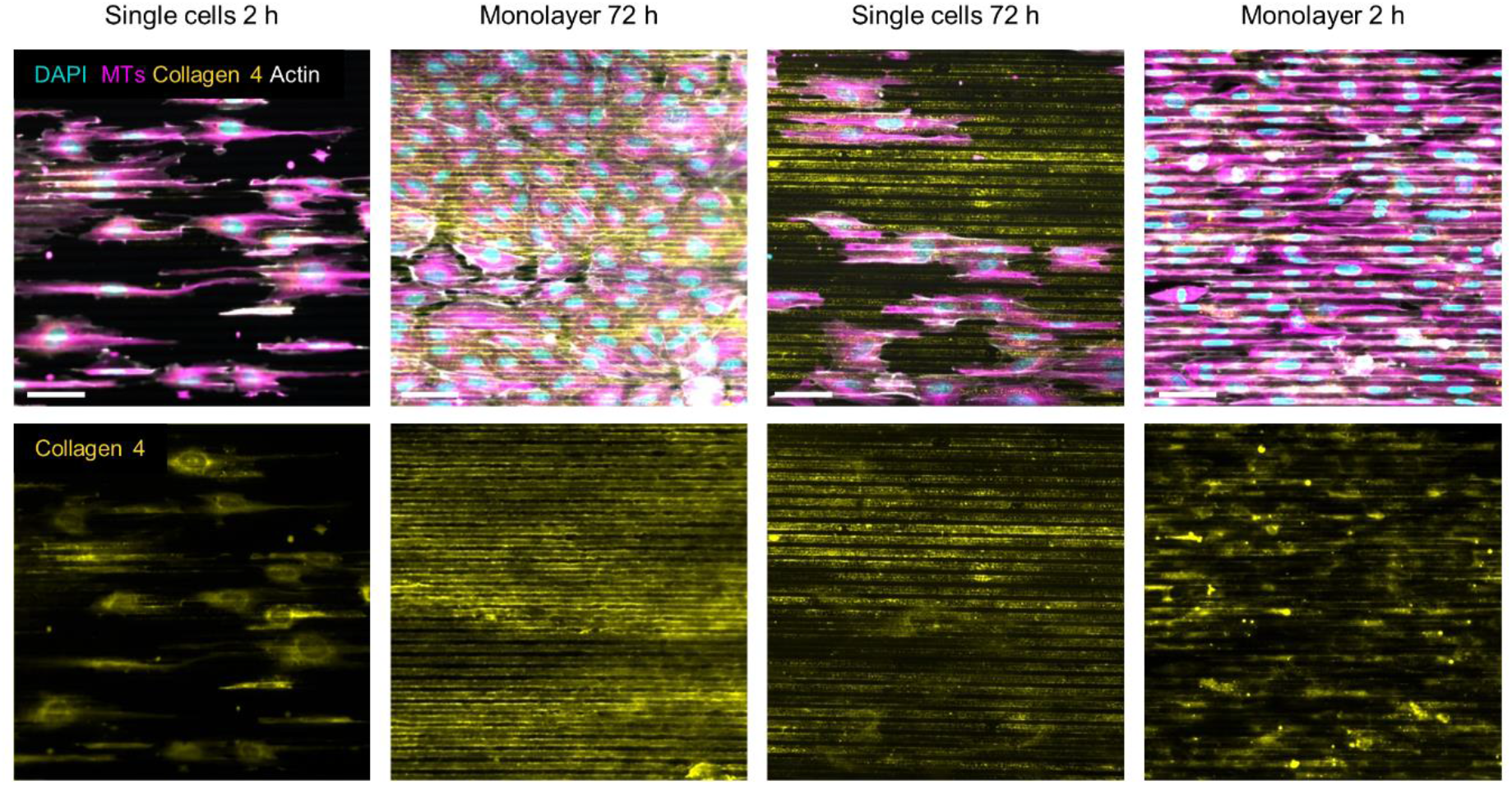
Influence of time vs. density on the secreted basement membrane. Immunostaining against DAPI (cyan), microtubules (α-tubulin, magenta), collagen 4 (yellow), and actin (grey) for single cells after 2 h of culture, monolayers after 72 h of culture, single cells after 72 h of culture, or monolayers after 2 h of culture. Scale bars, 50 μm.

**Supplementary video 1**: recording of Lifeact-Ruby HUVECs on 5 x 5 x 1 μm (w x s x d) grooves (horizontal). Scale bar, 20 μm.

**Supplementary video 2**: recording of Lifeact-Ruby HUVECs on 5 x 5 x 5 μm (w x s x d) grooves (horizontal). Scale bar, 20 μm.

## Notes

### Competing Interest Statement

The authors have declared no competing interest.

## References

Akanuma, T., Chen, C., Sato, T., Merks, R. M. H. and Sato, T. N. (2016). Memory of cell shape biases stochastic fate decision-making despite mitotic rounding. Nat. Commun. 7, 11963.

Andersson, A. S., Olsson, P., Lidberg, U. and Sutherland, D. (2003). The effects of continuous and discontinuous groove edges on cell shape and alignment. Exp. Cell Res. 288, 177–88.

Antonini, S., Meucci, S., Jacchetti, E., Klingauf, M., Beltram, F., Poulikakos, D., Cecchini, M. and Ferrari, A. (2015). Sub-micron lateral topography affects endothelial migration by modulation of focal adhesion dynamics. Biomed. Mater.

Atlas, Y., Gorin, C., Novais, A., Marchand, M. F., Chatzopoulou, E., Lesieur, J., Bascetin, R., Binet-Moussy, C., Sadoine, J., Lesage, M., et al. (2021). Microvascular maturation by mesenchymal stem cells in vitro improves blood perfusion in implanted tissue constructs. Biomaterials 268, 120594.

Bignon, M., Pichol-Thievend, C., Hardouin, J., Malbouyres, M., Bréchot, N., Nasciutti, L., Barret, A., Teillon, J., Guillon, E., Etienne, E., et al. (2011). Lysyl oxidase-like protein-2 regulates sprouting angiogenesis and type IV collagen assembly in the endothelial basement membrane. Blood 118, 3979–3989.

Curtis, A. S.. and Clark, P. (1990). The Effect of Topographic and Mechanical Properties of Materials on Cell Behaviour. Crit. Rev. Biocompat. 5, 343–362.

Dessalles, C. A., Leclech, C., Castagnino, A. and Barakat, A. I. (2021). Integration of substrate-and flow-derived stresses in endothelial cell mechanobiology. Commun. Biol. 4,.

Dewey, C. F., Bussolari, S. R., Gimbrone, M. A. and Davies, P. F. (1981). The Dynamic Response of Vascular Endothelial Cells to Fluid Shear Stress. J. Biomech. Eng. 103,.

Franco, D., Klingauf, M., Bednarzik, M., Cecchini, M., Kurtcuoglu, V., Gobrecht, J., Poulikakos, D. and Ferrari, A. (2011). Control of initial endothelial spreading by topographic activation of focal adhesion kinase. Soft Matter 7, 7313–7324.

Fujita, S., Ohshima, M. and Iwata, H. (2009). Time-lapse observation of cell alignment on nanogrooved patterns. J. R. Soc. Interface 6, S269–S277.

Hahn, C. and Schwartz, M. A. (2009). Mechanotransduction in vascular physiology and atherogenesis. Nat. Rev. Mol. Cell Biol. 10,.

Hayer, A., Shao, L., Chung, M., Joubert, L. M., Yang, H. W., Tsai, F. C., Bisaria, A., Betzig, E. and Meyer, T. (2016). Engulfed cadherin fingers are polarized junctional structures between collectively migrating endothelial cells. Nat. Cell Biol. 18, 1311–1323.

Helmlinger, G., Geiger, R. V., Schreck, S. and Nerem, R. M. (1991). Effects of Pulsatile Flow on Cultured Vascular Endothelial Cell Morphology. J. Biomech. Eng. 113,.

Hoelzle, M. K. and Svitkina, T. (2012). The cytoskeletal mechanisms of cell-cell junction formation in endothelial cells. Mol. Biol. Cell 23, 310–323.

Huveneers, S., Oldenburg, J., Spanjaard, E., van der Krogt, G., Grigoriev, I., Akhmanova, A., Rehmann, H. and de Rooij, J. (2012). Vinculin associates with endothelial VE-cadherin junctions to control force-dependent remodeling. J. Cell Biol. 196, 641–652.

Leclech, C. and Barakat, A. I. (2021). Is there a universal mechanism of cell alignment in response to substrate topography? Cytoskeleton.

Leclech, C. and Villard, C. (2020). Cellular and Subcellular Contact Guidance on Microfabricated Substrates. Front. Bioeng. Biotechnol. 8–551505.

Leclech, C., Natale, C. F. and Barakat, A. I. (2020). The basement membrane as a structured surface - role in vascular health and disease. J. Cell Sci. 133, jcs239889-undefined.

Lee, K., Kim, E. H., Oh, N., Tuan, N. A., Bae, N. H., Lee, S. J., Lee, K. G., Eom, C. Y., Yim, E. K. and Park, S. (2016). Contribution of actin filaments and microtubules to cell elongation and alignment depends on the grating depth of microgratings. J. Nanobiotechnology 14, 35.

Levesque, M. J., Liepsch, D., Moravec, S. and Nerem, R. M. (1986). Correlation of endothelial cell shape and wall shear stress in a stenosed dog aorta. Arterioscler. An Off. J. Am. Hear. Assoc. Inc. 6, 220–229.

Liliensiek, S. J., Wood, J. A., Yong, J., Auerbach, R., Nealey, P. F. and Murphy, C. J. (2010). Modulation of human vascular endothelial cell behaviors by nanotopographic cues. Biomaterials 31, 5418–5426.

Lou, H. Y., Zhao, W., Zeng, Y. and Cui, B. (2018). The Role of Membrane Curvature in Nanoscale Topography-Induced Intracellular Signaling. Acc. Chem. Res. 51, 1046–1053.

Lum, R. M., Wiley, L. M. and Barakat, A. I. (2000). Influence of different forms of fluid shear stress on vascular endothelial TGF-beta1 mRNA expression. Int. J. Mol. Med.

Luxenburg, C. and Zaidel-Bar, R. (2019). From cell shape to cell fate via the cytoskeleton — Insights from the epidermis. Exp. Cell Res. 378, 232–237.

Maruthamuthu, V., Sabass, B., Schwarz, U. S. and Gardel, M. L. (2011). Cell-ECM traction force modulates endogenous tension at cell-cell contacts. Proc. Natl. Acad. Sci. 108, 4708–4713.

Millán, J., Cain, R. J., Reglero-Real, N., Bigarella, C., Marcos-Ramiro, B., Fernández-Martín, L., Correas, I. and Ridley, A. J. (2010). Adherens junctions connect stress fibres between adjacent endothelial cells. BMC Biol. 8,.

Mooney, D., Hansen, L., Vacanti, J., Langer, R., Farmer, S. and Ingber, D. (1992). Switching from differentiation to growth in hepatocytes: Control by extracellular matrix. J. Cell. Physiol. 151, 497–505.

Morgan, J. T., Wood, J. A., Shah, N. M., Hughbanks, M. L., Russell, P., Barakat, A. I. and Murphy, C. J. (2012). Integration of basal topographic cues and apical shear stress in vascular endothelial cells. Biomaterials 33, 4126–35.

Natale, C. F., Lafaurie-Janvore, J., Ventre, M., Babataheri, A. and Barakat, A. I. (2019). Focal adhesion clustering drives endothelial cell morphology on patterned surfaces. J. R. Soc. Interface 16, 20190263.

Ngandu Mpoyi, E., Cantini, M., Sin, Y. Y., Fleming, L., Zhou, D. W., Costell, M., Lu, Y., Kadler, K., García, A. J., Van Agtmael, T., et al. (2020). Material-driven fibronectin assembly rescues matrix defects due to mutations in collagen IV in fibroblasts. Biomaterials 252, 120090.

Noethel, B., Ramms, L., Dreissen, G., Hoffmann, M., Springer, R., Rübsam, M., Ziegler, W. H., Niessen, C. M., Merkel, R. and Hoffmann, B. (2018). Transition of responsive mechanosensitive elements from focal adhesions to adherens junctions on epithelial differentiation. Mol. Biol. Cell 29, 2317–2325.

Ray, A., Lee, O., Win, Z., Edwards, R. M., Alford, P. W., Kim, D.-H. and Provenzano, P. P. (2017). Anisotropic forces from spatially constrained focal adhesions mediate contact guidance directed cell migration. Nat. Commun. 8, 4923.

Reinhart-King, C. A., Dembo, M. and Hammer, D. A. (2005). The dynamics and mechanics of endothelial cell spreading. Biophys. J. 89, 676–689.

Sahai, E. (2007). Illuminating the metastatic process. Nat. Rev. Cancer 7, 737–749.

Saito, A. C., Matsui, T. S., Ohishi, T., Sato, M. and Deguchi, S. (2014). Contact guidance of smooth muscle cells is associated with tension-mediated adhesion maturation. Exp. Cell Res. 327, 1–11.

Sales, A., Holle, A. W. and Kemkemer, R. (2017). Initial contact guidance during cell spreading is contractility-independent. Soft Matter 13, 5158–5167.

Sharma, V. P., Beaty, B. T., Patsialou, A., Liu, H., Clarke, M., Cox, D., Condeelis, J. S. and Eddy, R. J. (2012). Reconstitution of in vivo macrophage-tumor cell pairing and streaming motility on one-dimensional micro-patterned substrates. IntraVital 1, 77–85.

Singhvi, R., Kumar, A., Lopez, G. P., Stephanopoulos, G. N., Wang, D. I. C., Whitesides, G. M. and Ingber, D. E. (1994). Engineering Cell Shape and Function. Science (80-.). 264, 696–698.

Song, K. H., Kwon, K. W., Song, S., Suh, K. Y. and Doh, J. (2012). Dynamics of T cells on endothelial layers aligned by nanostructured surfaces. Biomaterials.

Sprague, E. A., Tio, F., Ahmed, S. H., Granada, J. F. and Bailey, S. R. (2012). Impact of parallel micro-engineered stent grooves on endothelial cell migration, proliferation, and function: An in vivo correlation study of the healing response in the coronary swine model. Circ. Cardiovasc. Interv. 5, 499–507.

Stefopoulos, G., Giampietro, C., Falk, V., Poulikakos, D. and Ferrari, A. (2017). Facile endothelium protection from TNF-α inflammatory insult with surface topography. Biomaterials 138, 131–141.

Stringer, C., Wang, T., Michaelos, M. and Pachitariu, M. (2021). Cellpose: a generalist algorithm for cellular segmentation. Nat. Methods 18, 100–106.

Tabdanov, E. D., Puram, V., Zhovmer, A. and Provenzano, P. P. (2018). Microtubule-Actomyosin Mechanical Cooperation during Contact Guidance Sensing. Cell Rep. 328–338.

Tan, C. H., Muhamad, N. and Abdullah, M. M. A. B. (2017). Surface Topographical Modification of Coronary Stent: A Review. In IOP Conference Series: Materials Science and Engineering, p. Institute of Physics Publishing.

Teixeira, A. I., Abrams, G. A., Bertics, P. J., Murphy, C. J. and Nealey, P. F. (2003). Epithelial contact guidance on well-defined micro-and nanostructured substrates. J. Cell Sci. 116, 1881–1892.

Umana-Diaz, C., Pichol-Thievend, C., Marchand, M. F., Atlas, Y., Salza, R., Malbouyres, M., Barret, A., Teillon, J., Ardidie-Robouant, C., Ruggiero, F., et al. (2020). Scavenger Receptor Cysteine-Rich domains of Lysyl Oxidase-Like2 regulate endothelial ECM and angiogenesis through non-catalytic scaffolding mechanisms. Matrix Biol. 88, 33–52.

Vesga, B., Hernandez, H., Higuera, S., Gasior, P., Echeveri, D., Delgado, J. A., Dager, A., Arana, C., Simonton, C., Maehara, A., et al. (2017). Biological effect of microengineered grooved stents on strut healing: A randomised OCT-based comparative study in humans. Open Hear. 4,.

Wang, Z., Liu, C., Xiao, Y., Gu, X., Xu, Y., Dong, N., Zhang, S., Qin, Q. and Wang, J. (2019). Remodeling of a Cell-Free Vascular Graft with Nanolamellar Intima into a Neovessel. ACS Nano 13, 10576–10586.

Weigelin, B., Bakker, G.-J. and Friedl, P. (2012). Intravital third harmonic generation microscopy of collective melanoma cell invasion. IntraVital 1,.

Zimmermann, J., Camley, B. A., Rappel, W.-J. and Levine, H. (2016). Contact inhibition of locomotion determines cell–cell and cell–substrate forces in tissues. Proc. Natl. Acad. Sci. 113, 2660–2665.

